# Astrocyte-derived MFG-E8 facilitates microglial synapse elimination in Alzheimer’s disease mouse models

**DOI:** 10.1101/2024.08.31.606944

**Authors:** Dimitra Sokolova, Shari Addington Ghansah, Francesca Puletti, Tatiana Georgiades, Sebastiaan De Schepper, Yongjing Zheng, Gerard Crowley, Ling Wu, Javier Rueda-Carrasco, Angeliki Koutsiouroumpa, Philip Muckett, Oliver J. Freeman, Baljit S. Khakh, Soyon Hong

## Abstract

Region-specific synapse loss is an early pathological hallmark in Alzheimer’s disease (AD). Emerging data in mice and humans highlight microglia, the brain-resident macrophages, as cellular mediators of synapse loss; however, the upstream modulators of microglia-synapse engulfment remain elusive. Here, we report a distinct subset of astrocytes, which are glial cells essential for maintaining synapse homeostasis, appearing in a region-specific manner with age and amyloidosis at onset of synapse loss. These astrocytes are distinguished by their peri-synaptic processes which are ‘bulbous’ in morphology, contain accumulated p62-immunoreactive bodies, and have reduced territorial domains, resulting in a decrease of astrocyte-synapse coverage. Using integrated *in vitro* and *in vivo* approaches, we show that astrocytes upregulate and secrete phagocytic modulator, milk fat globule-EGF factor 8 (MFG-E8), which is sufficient and necessary for promoting microglia-synapse engulfment in their local milieu. Finally, we show that knocking down *Mfge8* specifically from astrocytes using a viral CRISPR-saCas9 system prevents microglia-synapse engulfment and ameliorates synapse loss in two independent amyloidosis mouse models of AD. Altogether, our findings highlight astrocyte-microglia crosstalk in determining synapse fate in amyloid models and nominate astrocytic MFGE8 as a potential target to ameliorate synapse loss during the earliest stages of AD.

## Introduction

Region-specific vulnerability of synapses to loss and dysfunction is an early pathological hallmark and a significant correlate for cognitive decline in Alzheimer’s disease (AD)^1–6^. As the primary tissue-resident macrophages of brain parenchyma, microglia play a critical role in neuronal synapse loss in various neurologic diseases, including AD^7–16^. We and others have previously shown that microglia engulf synapses via the classical complement cascade (C1q/C3)^7,10,15^. However, what determines which synapses become vulnerable to microglial elimination is unclear. Astrocytes are sponge-shaped glial cells that intimately interact with synapses via peri-synaptic processes, which enables them to perform numerous functions critical for maintaining synapse homeostasis^17–20^. These functions include regulating the number of synapses in both health and disease^10,14,21–24^. Here, we report that astrocytes are responsible for creating a local niche that promotes microglial engulfment and loss of excitatory post-synaptic Homer1. Specifically, we found a subset of astrocytes characterized by spherical p62-immunoreactive ‘bulbous’ peri-synaptic processes appearing in a highly region-specific manner, including the hippocampus, at the onset of synapse loss in the hAPP NL-F (NL-F) knock-in (KI) mouse model of AD^25^. Mechanistically, we identified that these astrocytes upregulate and secrete milk fat globule-EGF factor 8 (MFG-E8), a known modulator of phagocytosis in their local milieu^14,26–28^. We further report that knockdown of MFG-E8 specifically from astrocytes prevents Homer1 engulfment by microglia and ameliorates excitatory synapse loss in multiple models of amyloidosis. Our findings demonstrate a novel local neighborhood concept for astrocyte-neuron-microglia network in modulating synapse loss in AD.

## Results

### A morphologically distinct p62-accumulated astrocyte subset increases with age and amyloidosis in the mouse hippocampus

Astrocytes have elaborate processes that intimately enwrap synapses and provide critical trophic support to maintain synaptic health and function^17–20^. This intimate apposition of astrocytic processes to synapses is critical for assessing proper astrocyte-neuron interactions. However, whether the full morphological complexity of astrocytes and their astrocyte-synapse coverages change in early stages of AD and if so, whether these alterations have implications for synapse loss, are poorly understood. To address this, we assessed the morphology of astrocytes and their processes in the hippocampus of a slow-progressing second-generation mouse model of AD, the hAPP NL-F (NL-F) knock-in (KI)^25^, where humanized familial AD mutations are under the control of the endogenous APP promoter. In the NL-F KI model, where age is a critical factor of amyloidosis, astrocyte calcium dysfunction^29^ and microglial engulfment of synapses^8,9^ are observed at 3- and 6-months (mo) of age respectively in the hippocampus, prior to robust amyloid-β (Aβ) plaque accumulation, which is not observed until 12-15-mo^25^.

Hippocampal astrocytes have a complex, dense, and bushy morphology, which is critical for their function and interaction with synapses^20^. However, fine peri-synaptic astrocytic processes cannot be labeled by glial fibrillary acidic protein (GFAP) immunostaining **(Fig S1A)**^30–32^. To visualize the complexity of astrocytic processes *in vivo*, we injected AAV2/5-*GfaABC_1_D*-tdTomato (tdTomato) virus^33^ into the hippocampus, which selectively and fully labels astrocytes and their processes **(Fig 1A)**. We found that the 6-mo NL-F KI hippocampal tdTomato-labeled astrocytes have a significantly smaller surface area than their wild-type (WT) counterparts **(Figs 1B** and **C)**^34^. Notably, within the NL-F KI hippocampus, we identified a morphologically distinct subset of astrocytes with deformed spherical peri-synaptic processes, from here on referred to as ‘bulbous’ **(Fig 1D)**. We found that astrocytes with bulbous processes have a significantly reduced territory volume as compared to their neighboring astrocytes with bushy processes within the same hippocampal brain region, as measured by high-resolution confocal microscopy and Imaris 3D surface rendering **(Fig 1** **E).** Using immunohistochemistry (IHC) with S100 calcium-binding protein B (S100β), which can capture complex astrocytic processes^35^, we observed similar bulbous processes in non-tdTomato injected NL-F KI brains, confirming that this phenotype is not a viral artefact **(Fig S1A)**. In contrast, IHC with GFAP, which only labels primary astrocytic branches^30,31^, was insufficient to label astrocytic peri-synaptic processes **(Fig S1A)** nor capture the changes in astrocyte volume between WT and NL-F KI mice **(Fig S1B)**.

**Fig 1.**
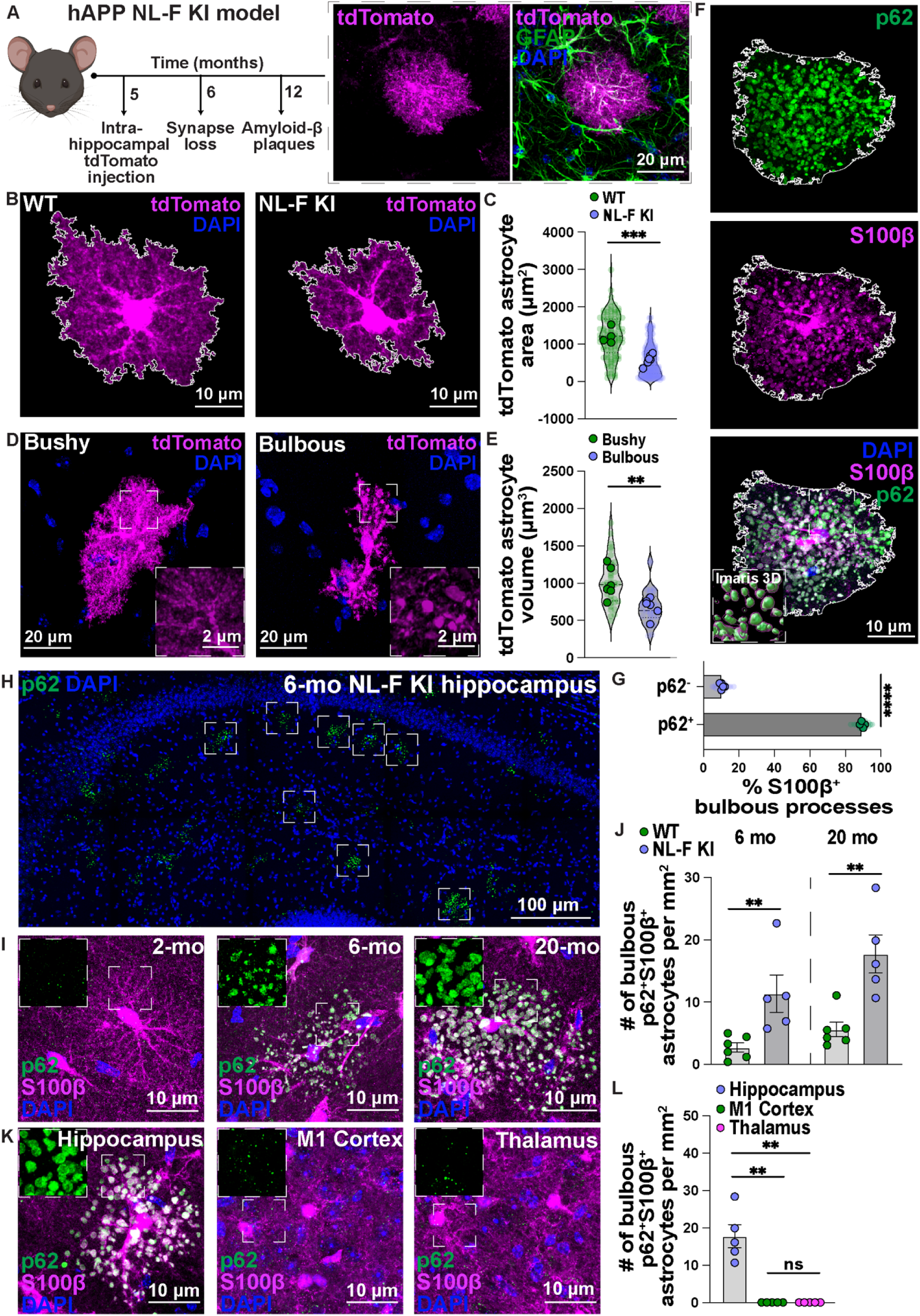
Identification of a unique p62-accumulated bulbous astrocyte subset in the NL-F KI hippocampus. **A)** Schematic of NL-F KI mouse model and viral tdTomato labeling strategy to visualize complex hippocampal astrocyte processes. Immunostaining for GFAP (green) post intra-hippocampal AAV2/5-*GfaABC1D*-tdTomato (tdTomato) (magenta) injection. Scale bar = 10 μm. **B)** Representative images of tdTomato-labeled astrocytes (magenta) in the 6-mo WT and NL-F KI hippocampus. Scale bar = 10 μm. Images are modified on ImageJ as shown by white ROI. **C)** Quantification of the tdTomato labeled astrocyte area (μm^2^) using ImageJ. Transparent points = individual astrocytes (16-58 astrocytes per mouse; 104 WT and 183 NL-F KI astrocytes sampled in total); full points = mouse average of ROIs (n=4-6 male mice per genotype). Unpaired two-tailed student’s t-test on mouse average. **D)** Representative images of tdTomato-labeled (magenta) astrocytes in the 6-mo hippocampus showing a bushy and bulbous astrocyte. Scale bar = 20 μm. Insets show representative zoom-ins of processes. Scale bar = 2 μm. **E)** Quantification of the tdTomato-labeled astrocytic volume (μm^3^) using Imaris 3D surface rendering. Transparent points = individual astrocytes (5-7 astrocytes per condition; 35 NL-F KI bushy and 31 NL-F KI bulbous astrocytes sampled in total); full points = mouse average of ROIs (n=6 male mice). Paired two-tailed student’s t-test on mouse average. **F)** Representative image of p62 (green) and S100β (magenta) immunoreactivity in the hippocampus. Scale bar = 10 μm. Images are modified on ImageJ as shown by white ROI. Inset shows representative Imaris 3D reconstruction of p62 within the S100β^+^ astrocytic surface. **G)** Quantification of the % colocalization between p62 and S100β^+^ bulbous astrocytes in the hippocampus using ImageJ. Transparent points = individual astrocytes (3 astrocytes per mouse; 15 astrocytes sampled in total per condition), full points = mouse average of ROIs (n=5 male mice). Ratio paired two-tailed student’s t-test on mouse average. **H)** Representative image of p62 (green) immunoreactivity in the 6-mo NL-F KI hippocampus. Scale bar = 400 μm. Insets highlight some regions with bulbous astrocytes. **I)** Representative images of p62 (green) and S100β (magenta) immunoreactivity in the 2-, 6- and 20-mo hippocampus. Scale bar = 10 μm. Insets show representative zoom-ins of p62-immunostaining. **J)** Quantification of the number of p62^+^ S100β^+^ bulbous astrocytes per mm^2^ in the hippocampus in 6-mo (left) and 20-mo (right). Each point = 1 mouse (n=5-6 male mice). Two-tailed Mann-Whitney (6-mo) or unpaired two-tailed student’s t-test (20-mo) on mouse average. Dashed line represents graph split. **K)** Representative images of p62 (green) and S100β (magenta) immunoreactivity in the hippocampus, motor cortex (M1) and thalamus. Scale bar = 10 μm. Insets show representative zoom-ins of p62-immunostaining. **L)** Quantification of the number of p62^+^ S100β^+^ bulbous astrocytes per mm^2^ in hippocampus, motor cortex (M1) and thalamus. Each point = 1 mouse (n=5 male mice). Kruskal-Wallis test (p<0.001) followed by the Dunn’s multiple comparisons test on mouse average. All data shown as mean ± SEM. p-values shown ns P>0.05; *P<0.05; **P<0.01; ***P<0.001; ****P<0.0001.

Morphologically, the bulbous astrocytic processes resembled structures identifiable by their labeling with selective autophagy cargo receptor p62/SQSTM1 (p62), known as Lafora bodies, polyglycosan bodies and corpora amylacea, which have been reported in mouse and human tissues in various contexts including AD^36–42^. Indeed, by IHC, we found a striking 90% colocalization between bulbous S100β^+^ processes and p62-immunoreactive spherical bodies **(Figs 1F** and **G)**. To investigate when bulbous astrocytes start appearing in the NL-F KI hippocampus, we examined three timepoints, i.e., 2-, 6- and 20-mo, reflecting pre-pathology, onset of synapse loss, and Aβ plaque-enriched time points, respectively **(Figs 1H-J)**^7–9,25^. There was no detectable level of the bulbous astrocytes at 2-mo in either genotype, but the bulbous astrocytes increased with age in both WT and NL-F KI hippocampus **(Figs S1C** and **D)**. At 6-mo, a key timepoint of synapse loss, levels of bulbous astrocytes were significantly higher in the NL-F KI hippocampus compared to those in the WT **(Fig 1J)**; we also observed higher levels in the Aβ plaque-laden 20-mo NL-F KI hippocampus vs. age- and sex-matched WT controls. These data suggest that amyloidosis exacerbates the formation of the bulbous astrocytic processes. Interestingly, we found little spatial association between p62-accumulated bulbous astrocytes and 6E10-immunoreactive Aβ aggregated deposits (60 μm median proximity distance) in the Aβ plaque-laden 20-mo NL-F KI hippocampus **(Figs S1E-G)**, suggesting that these astrocytes are not a peri-plaque phenotype^43^. Next, we explored the spatial distribution of the bulbous astrocytes across different brain regions. p62-accumulated S100β^+^ bulbous astrocytes were found in the hippocampus, a brain region affected in AD^44,45^, but not in the primary motor cortex or the thalamus **(Figs 1K and L)**. We further performed serial cross-section imaging across the cerebrum, which showed presence of p62-accumulated S100β astrocytes in distinct brain regions, i.e., piriform cortex, hippocampus and subiculum **(Fig S2)**^38^. Finally, we found round p62^+^ S100β^+^ ALDH1L1^+^ astrocytic processes by IHC in the hippocampus of post-mortem control and AD human tissue **(Fig S3)**, in line with previous reports^36,46,47^, suggesting that analogous astrocytic phenotypes may exist in both mouse and human hippocampi.

### Decreased astrocyte-synapse coverage in the pre-plaque NL-F KI hippocampus

The spatiotemporal emergence of bulbous peri-synaptic astrocytic processes and synapse loss in the NL-F KI hippocampus raised the question of whether there is a change in the astrocyte-synapse associations in the NL-F KI model. To that end, we used super-resolution Airyscan confocal microscopy to assess the colocalization between excitatory synaptic puncta, as defined by colocalized or closely apposed Synaptophysin- and Homer1-immunoreactive pre- and post-synaptic puncta, respectively, and the excitatory amino acid transporter 1 (EAAT1), an astrocytic glutamate transporter localized on astrocytic membranes in synaptically dense regions **(Figs 2A-F)**^48,49^. Consistent with previous findings, there was a significant decrease in the density of excitatory synapses **(Fig 2B)**^8,9^ as well as a decrease in the number of EAAT1-immunoreactive puncta **(Fig 2C)**^50–52^ in the hippocampal CA1 stratum radiatum (SR) of 6-mo NL-F KI mice as compared to those of WT mice. Interestingly, there was a significantly lower percentage of synapses colocalized with astrocytic EAAT1 as well as a decreased proportion of EAAT1 associated with excitatory synapses in the NL-F KI hippocampal CA1 SR compared to WT controls **(Figs 2D** and **E)**, suggesting that there is a reduction in astrocyte-synapse association in the 6-mo NL-F KI hippocampus.

**Fig 2.**
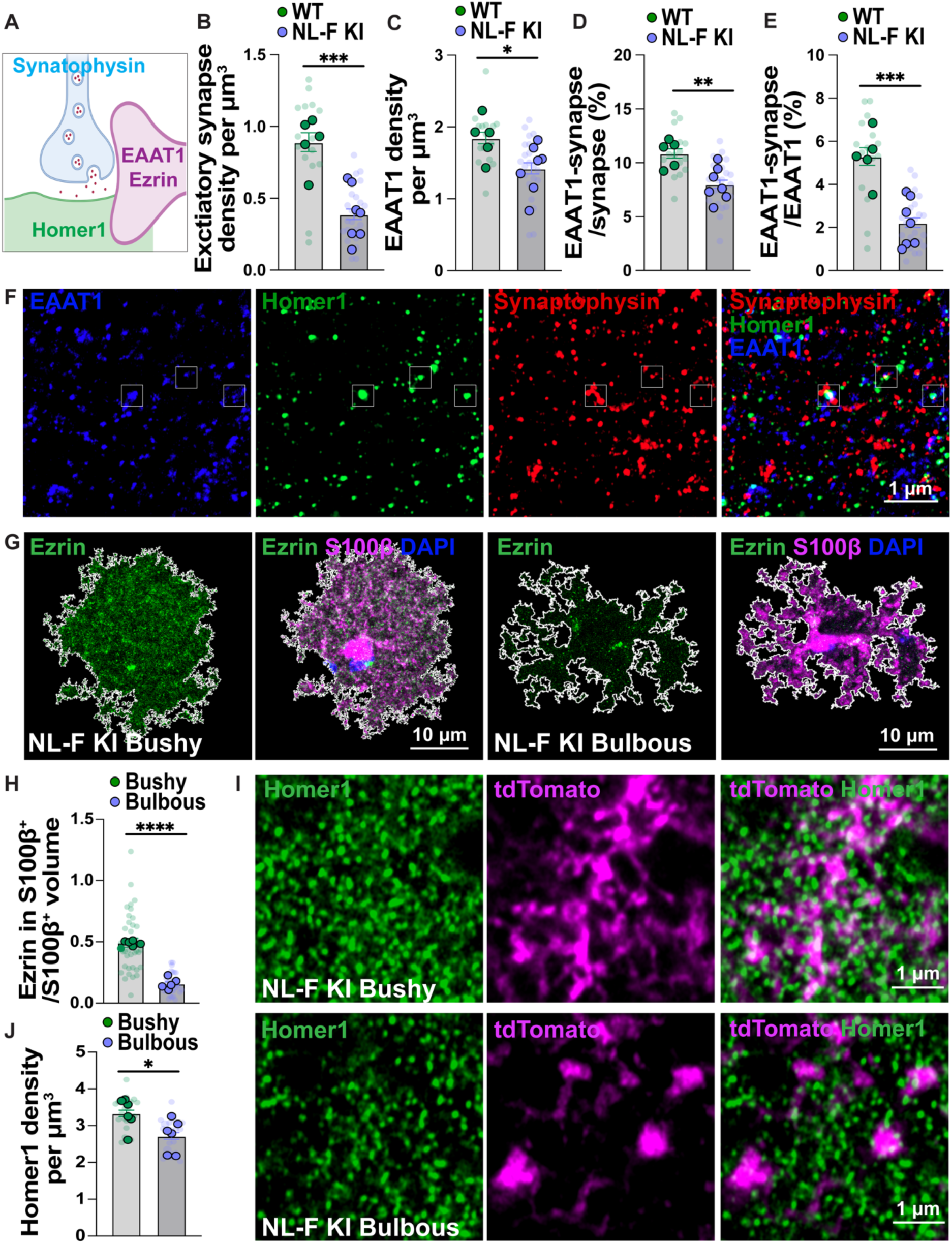
Loss of astrocyte-synapse interactions and increased loss of excitatory post-synaptic marker Homer1 near hippocampal NL-F KI bulbous astrocytes. **A)** Schematic of astrocytic-synapse interactions. **B-F)** Representative images and quantification of astrocytic EAAT1-, excitatory post-synaptic Homer1- and pre-synaptic Synaptophysin-immunoreactive puncta in the 6-mo hippocampal CA1 SR using super-resolution Airyscan confocal microscopy. **B)** Quantification of the number of colocalized Homer1- and Synaptophysin-immunoreactive spots shown as density per µm^3^ using Imaris. Transparent points = individual ROIs (3 ROIs per mouse; 15 WT and 21 NL-F KI ROIs sampled in total); full points = mouse average of ROIs (n=5-7 male mice per genotype). Unpaired two-tailed student’s t-test on mouse average. **C)** Quantification of the number of EAAT1-immunoreactive spots shown as density per µm^3^ using Imaris. Transparent points = individual ROIs (3 ROIs per mouse; 15 WT and 21 NL-F KI ROIs sampled in total); full points = mouse average of ROIs (n=5-7 male mice per genotype). Unpaired two-tailed student’s t-test on mouse average. **D)** Quantification of the number of colocalized EAAT1, Homer1 and Synaptophysin spots divided by the number of colocalized Homer1 and Synaptophysin spots shown as %. Transparent points = individual ROIs (3 ROIs per mouse; 15 WT and 21 NL-F KI ROIs sampled in total); full points = mouse average of ROIs (n=5-7 male mice per genotype). Unpaired two-tailed student’s t-test on mouse average. **E)** Quantification of the number of colocalized EAAT1, Homer1 and Synaptophysin spots divided by the number of EAAT1 spots using Imaris shown as %. Transparent points = individual ROIs (3 ROIs per mouse; 15 WT and 21 NL-F KI ROIs sampled in total); full points = mouse average of ROIs (n=5-7 male mice per genotype). Unpaired two-tailed student’s t-test on mouse average**. F)** Representative image of **B-E** for EAAT1 (blue), Homer1 (green) and Synaptophysin (red). Scale bar = 1 μm. Insets show regions of triple colocalization. **G)** Representative images of Ezrin (green) immunoreactivity in the 6-mo NL-F KI hippocampus near bushy or bulbous S100β^+^ astrocytes (magenta). Scale bar = 10 μm. Images are modified on ImageJ as shown by white ROI. **H)** Quantification of the total Ezrin spot volume on S100β^+^ astrocytes/S100β volume using Imaris 3D surface rendering and surface-spot colocalization. Transparent points = individual astrocytes (5-8 astrocytes per condition; 36 NL-F KI bushy and 30 NL-F KI bulbous astrocytes sampled in total); full points = mouse average of ROIs (n=5 male mice). Paired two-tailed student’s t-test on mouse average. **I)** Representative images for excitatory post-synaptic Homer1-immunoreactive puncta (green) in the 6-mo NL-F KI hippocampus near bushy or bulbous S100β^+^ astrocytes (magenta) using super-resolution Airyscan confocal microscopy. Scale bar = 1 μm. **J)** Quantification of the number of Homer1-immunoreactive puncta shown as density per µm^3^ using Imaris. Transparent points = individual ROIs (3 ROIs per condition; 18 ROIs sampled in total per condition), full points = mouse average of ROIs (n=6 male mice). Wilcoxon matched pairs signed rank test on mouse average. All data shown as mean ± SEM. p-values shown ns P>0.05; *P<0.05; **P<0.01; ***P<0.001; ****P<0.0001.

We further examined for the levels of Ezrin, an integral leaflet-structural protein critical for astrocyte morphology and synapse coverage^34,48,53^, within S100β-immunoreactive astrocytes. We assessed for Ezrin levels in astrocytes with bulbous peri-synaptic processes vs. bushy ones using high-resolution confocal microscopy and Imaris 3D surface rendering. We found that the NL-F KI hippocampal astrocytes with bulbous peri-synaptic processes contained significantly less Ezrin vs. those with bushy processes **(Figs 2G** and **H)**. In line with the decreased Ezrin and astrocyte-synaptic associations, we found a significantly decreased number of Homer1-immunoreactive synaptic puncta in the vicinity (< 40 μm radius) of bulbous astrocytic processes as compared to the vicinity of bushy astrocytic processes within the same NL-F KI CA1 SR hippocampus **(Figs 2I** and **J)**. These data altogether suggest a reduced synaptic coverage by the bulbous astrocytes and resulting synapse loss in their local milieu.

### Increased microglia-synapse engulfment in the local milieu of bulbous astrocytes in the NL-F KI hippocampus

Astrocytes, like microglia, have been shown to engulf synapses in various contexts including developmental synaptic pruning and in mouse models of AD pathology such as the TauP301S and APP/PS1^10,14,21–23^. Thus, we asked whether the bulbous astrocytes are engulfing synapses in the 6-mo NL-F KI CA1 SR hippocampus. Using high-resolution confocal microscopy and 3D Imaris surface rendering, we assessed the levels of post-synaptic Homer1 inside Lamp1^+^ lysosomes of astrocytes in 6-mo NL-F KI vs. WT hippocampus. To our surprise, we did not find any appreciable differences between NL-F KI vs. WT hippocampus in the degree of internalized Homer1 puncta inside astrocytic Lamp1^+^ lysosomes; in both assessments of GFAP-immunoreactive astrocytes **(Figs 3A** and **B**) and tdTomato-labeled astrocytes (**Figs 3C** and **D**). Close examination of bushy vs. bulbous astrocytes did not show any significant difference in the levels of Homer1-immunoreactive puncta inside astrocytic Lamp1 lysosomes between these two morphologically distinct astrocytes in the NLF KI brain **(Figs 3C** and **D)**. These data altogether suggest that astrocytes are not engulfing Homer1 synapses, at least not during this early pre-plaque stage in the NL-F KI hippocampus.

**Fig 3.**
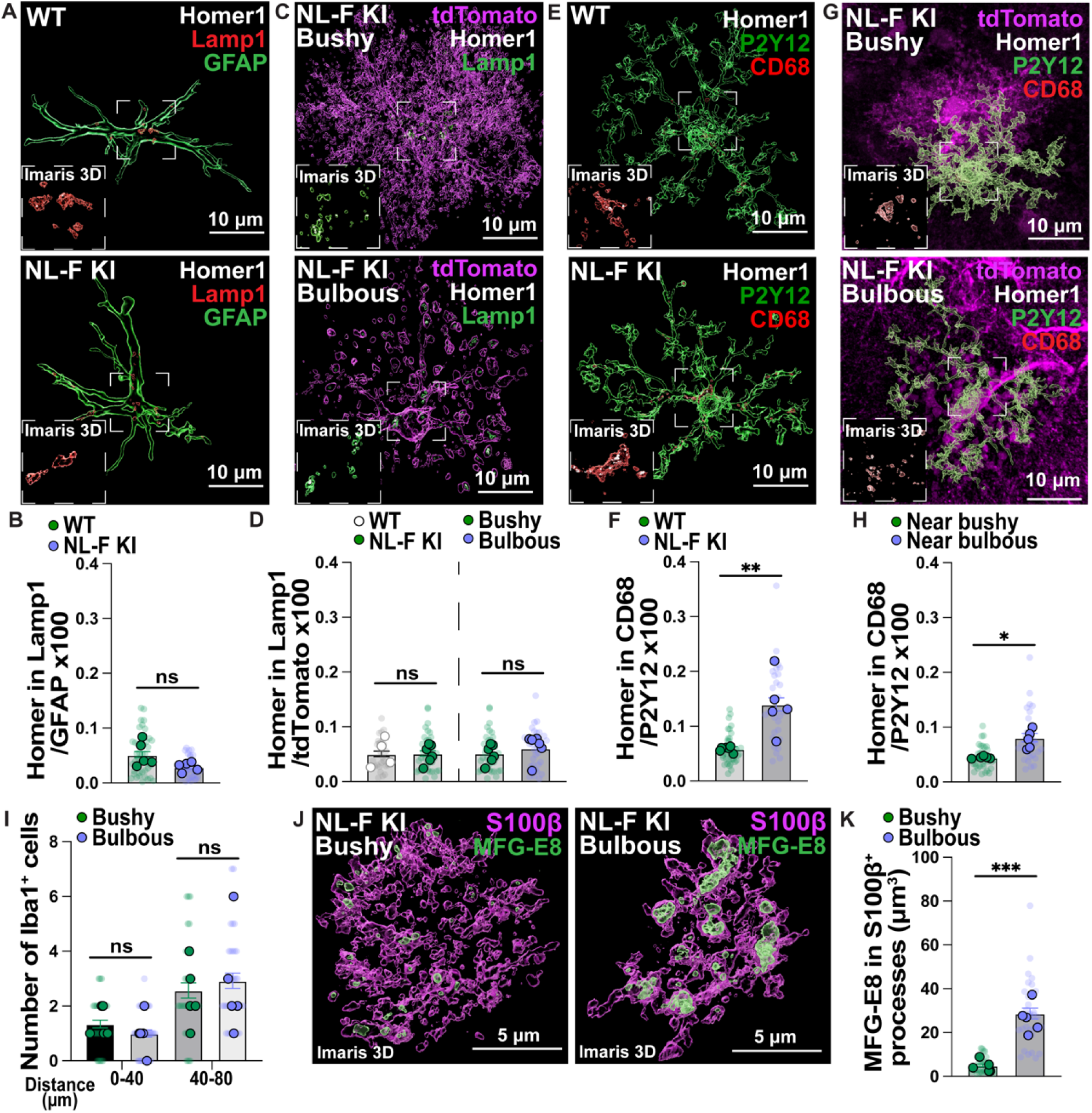
Increased microglia-Homer1 engulfment near hippocampal NL-F KI bulbous astrocytes. **A)** Representative 3D rendered images of excitatory post-synaptic Homer1 (white), Lamp1 lysosomes (red) and GFAP (green) in the 6-mo hippocampal CA1 SR. Scale bar = 10 μm. Inset shows representative zoom of Homer1 inside Lamp1^+^ astrocytic lysosomes. **B)** Quantification of astrocytic Homer1 engulfment using Imaris 3D surface rendering shown as: Homer1 volume in Lamp1^+^ lysosomes in GFAP^+^ astrocytic surface/GFAP volume x100. Transparent points = individual astrocytes (5-10 astrocytes per mouse; 40 WT and 39 NL-F KI astrocytes sampled in total); full points = mouse average of ROIs (n=5 male mice per genotype). Unpaired two-tailed student’s t-test with Welch’s correction on mouse average. **C)** Representative 3D rendered images of excitatory post-synaptic Homer1 (white), Lamp1 lysosomes (green) and tdTomato (magenta) in the 6-mo hippocampal CA1 SR. Scale bar = 10 μm. Inset shows representative zoom of Homer1 inside Lamp1^+^ astrocytic lysosomes. **D)** Quantification of astrocytic Homer1 engulfment using Imaris 3D surface rendering shown as: Homer1 volume in Lamp1^+^ lysosomes in tdTomato^+^ astrocytic surface/tdTomato volume x100. Transparent points = individual astrocytes (4-8 astrocytes per condition; 26 WT, 35 NL-F KI bushy and 31 NL-F KI bulbous astrocytes sampled in total), full points = mouse average of ROIs (n=4-6 male mice). Unpaired (WT vs NL-F KI) or paired (NL-F KI bushy vs NL-F KI bulbous) two-tailed student’s t-test on mouse average. Dashed line represents graph split. **E)** Representative 3D rendered images of excitatory post-synaptic Homer1 (white), CD68 lysosomes (red) and P2Y12 (green) in the 6-mo hippocampal CA1 SR. Scale bar = 10 μm. Inset shows representative zoom of Homer1 inside CD68^+^ microglial lysosomes. **F)** Quantification of microglial Homer1 engulfment using Imaris 3D surface rendering shown as: Homer1 volume in CD68^+^ lysosomes in P2Y12^+^ microglial surface/P2Y12 volume x100. Transparent points = individual microglia (5-6 microglia per mouse; 36 WT and 28 NL-F KI microglia sampled in total); full points = mouse average of ROIs (n=5-6 male mice per genotype). Two-tailed Mann Whitney test on mouse average. **G)** Representative 3D rendered images of excitatory post-synaptic Homer1 (white), CD68 lysosomes (red) and P2Y12 (green) in the 6-mo hippocampal CA1 SR near bushy or bulbous tdTomato^+^ (magenta) astrocytes. Scale bar = 10 μm. Inset shows representative zoom of Homer1 inside CD68^+^ microglial lysosomes. **H)** Quantification of microglial Homer1 engulfment using Imaris 3D surface rendering shown as: Homer1 volume in CD68^+^ lysosomes in P2Y12^+^ microglial surface/P2Y12 volume x100. Transparent point = individual microglia (5-7 microglia per condition; 32 and 28 microglia sampled in total near (< 5 μm away) bushy and bulbous NL-F KI astrocytes respectively), full points = mouse average of ROIs (n=6 male mice). Paired two-tailed student’s t-test on mouse average. **I)** Quantification of the number of Iba1^+^ microglia near bushy or bulbous NL-F KI astrocytes. Transparent points = individual astrocyte regions (4-10 regions per condition; 30 NL-F KI bushy and 33 NL-F KI bulbous astrocyte regions sampled in total); full points = mouse average of ROIs (n=5 male mice). Wilcoxon matched-pairs signed rank test followed by a Bonferroni-Dunn correction on mouse average. **J)** Representative 3D rendered images of MFG-E8 (green) inside S100β^+^ (magenta) bushy or bulbous astrocytic processes in the 6-mo NL-F KI hippocampus. Scale bar = 5 μm. **K)** Quantification of the volume of MFG-E8 within bushy or bulbous S100β^+^ astrocytic processes per μm^3^ using Imaris 3D surface rendering. Transparent points = individual astrocytes (3-5 astrocytes per condition; 27 NL-F KI bushy and 30 NL-F KI bulbous astrocytes sampled in total); full points = mouse average of ROIs (n=5 male mice). Paired two-tailed student’s t-test on mouse average. All data shown as mean ± SEM. p-values shown ns P>0.05; *P<0.05; **P<0.01; ***P<0.001; ****P<0.0001.

Next, we asked whether the loss of Homer1 near bulbous astrocytic processes is attributed to increased phagocytic engulfment by microglia in the local milieu. Microglia have been shown by us and others to engulf synapses in mouse models of Aβ pathology including in the NL-F KI model^7–10,15^. In accordance with earlier reports^7–10,15^, we found higher levels of Homer1-immunoreactive post-synaptic puncta inside CD68^+^ lysosomes of P2Y12^+^ microglial in the 6-mo NL-F KI vs. WT CA1 SR hippocampus **(Figs 3E** and **F)**. Notably, in the NL-F KI hippocampus, there was a significantly higher level of microglial engulfed Homer1 in the immediate (< 5 μm proximity) vicinity of tdTomato-labeled bulbous astrocytes as compared to that of the bushy astrocytes **(Figs 3G** and **H)**. There were no appreciable differences in the number of Iba1^+^ microglia between the vicinity of bulbous versus that of bushy tdTomato-labeled astrocytes **(Fig 3I),** suggesting that there is no recruitment of microglia *per se* but an enhanced trigger of microglia-synapse engulfment in bulbous local milieu. Altogether, these data suggest that the astrocytes themselves do not directly engulf Homer1 synapses at this early stage of amyloidosis but that there is an altered microenvironment near the bulbous peri-synaptic astrocytic processes that promotes microglia to engulf synapses.

### MFG-E8 in astrocyte-conditioned media is sufficient to promote microglial synaptosome engulfment

We next questioned whether the bulbous astrocytes, instead of directly engulfing synapses, may secrete factors to trigger microglia-synapse engulfment. Astrocytes possess a specialized secretome tailored to facilitate clearance^23,54,55^, some of which have been implicated in synapse elimination^10,14,21–24^. We first performed a reductionist *in vitro* microglial-synaptosome engulfment assay, where we tested the sufficiency of astrocyte-secreted factors in promoting microglia-synapse engulfment. Microglia were supplemented with astrocyte conditioned media (ACM) from primary astrocytes and live cell imaging was performed to measure the uptake of pHrodo-conjugated synaptosomes by microglia **(Fig S4**). We found that ACM significantly enhanced microglial engulfment of pHrodo-conjugated synaptosomes compared to microglia that did not receive any ACM **(Figs S4D** and **E**). To identify a candidate secreted phagocytic modulator in the ACM, we mined through existing bulk and single-cell RNA sequencing (scRNAseq) data sets^42,54–58^. Interestingly, a study highlighted the upregulation of secreted glycoprotein milk fat globule-EGF factor 8 (MFG-E8) in morphologically similar p62-accumulated astrocytes^42^.This is of high interest as MFG-E8 in the peripheral immune system has been shown to act as a secreted factor to promote engulfment by professional phagocytes^26^; dysregulation of MFG-E8 has also been reported in AD mouse and patient tissue^59–62^. Thus, we questioned whether astrocytes secrete MFG-E8 to promote microglia-synapse engulfment. Indeed, using Imaris 3D surface rendering, we found significantly higher MFG-E8-immunoreactivity in the local milieu of S100β-immunoreactive bulbous astrocytes in the NL-F KI hippocampus as compared to the bushy counterparts **(Figs 3J** and **K)**. To directly test the role of MFG-E8 in the *in vitro* engulfment assay, we immunodepleted (ID) MFG-E8 from the WT ACM, which contained physiological levels (ng/mL) of secreted MFG-E8 as shown by ELISA **(Fig S4F-H)**. Unlike the WT ACM, MFG-E8 ID ACM did not induce an increase in microglia synaptosome engulfment **(Figs S4D** and **E**); moreover, ACM from *Mfge8* knock-out (KO) primary astrocytes similarly failed to induce an increase **(Figs S4D** and **E**). Altogether, these data suggest that MFG-E8 secreted by astrocytes is sufficient to promote microglial engulfment of synaptosomes *in vitro*.

### Astrocytes are the main producers of *Mfge8* in the mouse and human brain parenchyma

In peripheral tissues such as the splenic germinal centers, macrophages produce MFG-E8 to facilitate engulfment^26,63^. Thus, MFG-E8 was assumed to be produced by microglia and has largely been studied *in vitro* in microglia^27,28,61^. However, multiple scRNAseq studies suggest that astrocytes, not microglia, express *Mfge8* in brain parenchyma^56–58,64^. To assess which cell types express *Mfge8* in the brain, we used single-molecule fluorescent *in situ* hybridization (smFISH; RNAScope) and observed that 90% of the *Mfge8* mRNA puncta are found on *Aldh1l1*^+^DAPI^+^ astrocytic nuclei but not *Cx3cr1*^+^DAPI^+^ microglial nuclei in the hippocampus **(Figs 4A** and **B)**. In line, RT-qPCR on fluorescence-activated cell sorting (FACS)-isolated cells showed *Mfge8* in sorted ACSA-2^+^CD45^-^ astrocytes but not CD45^+^CD11b^+^CX3CR1^+^ microglia **(Figs 4C; S5)**. Astrocyte-specific expression was also evident on protein level where MFG-E8 immunostaining colocalized with GFAP^+^ and S100β^+^ astrocytes but not with P2Y12^+^ microglia nor with NeuN^+^ neurons **(Figs S6A-C)**. Similarly, in a small cohort of post-mortem human patient tissue, we found that MFG-E8 immunostaining colocalizes with GFAP^+^ astrocytes in the frontal cortex **(Figs 4D; S6D),** in line with recent reports^60^. Of note, at the vasculature, *Mfge8* is also expressed by mural cells including smooth muscle cells **(Fig S6F)**^60^. Altogether, these results altogether confirm that astrocytes, not microglia, express *Mfge8* in the brain parenchyma **(Fig S6E)**.

**Fig 4.**
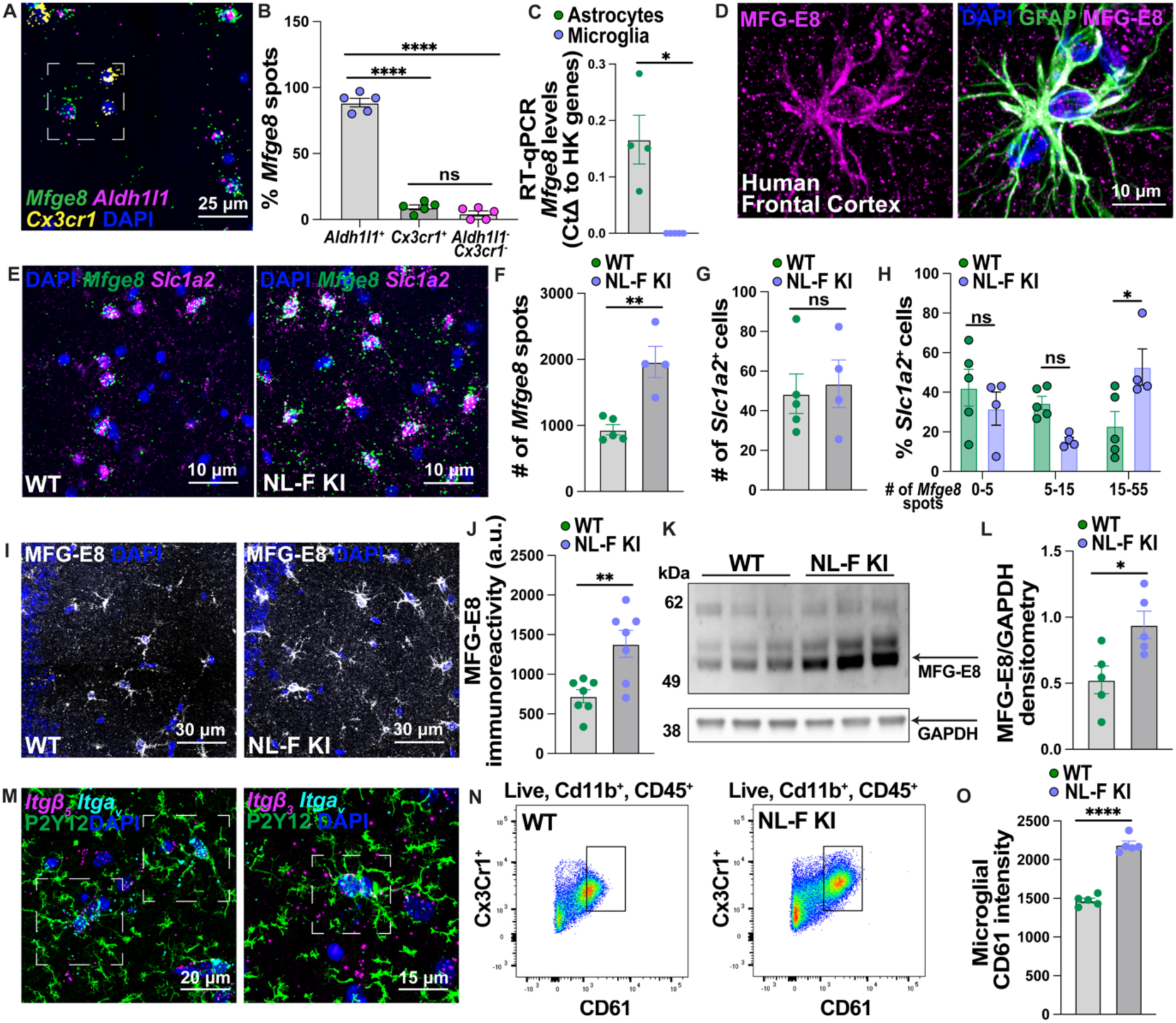
Increased astrocytic MFG-E8 signaling at the onset of synapse loss in the NL-F KI hippocampus. **A)** Representative image of RNAScope probing for *Mfge8* (green), *Aldh1l1^+^* astrocytes (magenta) and *Cx3cr1^+^* myeloid cells (yellow) in the hippocampus. Scale bar = 25 μm. Inset shows exemplary ROI. **B)** Quantification of the number of *Mfge8* mRNA spots on *Aldh1l1^+^* astrocytes, *Cx3cr1^+^* myeloid cells or *Aldh1l1^-^Cx3cr1^-^* cells in the hippocampus using Imaris. Data shown as a percentage of the total number of *Mfge8* spots in a large area of interest. Each point = 1 mouse (n=5 female mice). One-way ANOVA (F (2, 12) = 378.9, p<0.0001) followed by Bonferroni’s multiple comparisons post-hoc test on mouse average. **C)** RT-qPCR probing for *Mfge8* expression in FACS sorted ACSA2^+^CD45^-^ astrocytes and CX3CR1^+^CD45^+^CD11b^+^ microglia. Gene expression is normalized to the geomean of 3 house-keeping genes (*Actb*, *Gapdh* and *Rpl32*) by the Delta CT method. Each point = 1 mouse (n=3-4 female mice). Unpaired two-tailed student’s t-test with a Welch’s correction on mouse average. **D)** Representative images of MFG-E8 (magenta) and GFAP (green) immunoreactivity in the post-mortem human frontal cortex. Scale bar = 10 μm. **E)** RNAScope probing for *Mfge8* (green) on *Slc1a2^+^* (magenta) astrocytes in the 6-mo WT and NL-F KI hippocampus. Scale bar = 15 μm. **F)** Quantification of the total number of *Mfge8* spots in a large region of interest using Imaris. Each point = 1 mouse (n=4-5 mixed male and female mice per genotype). Unpaired two-tailed student’s t-test on mouse average. **G)** Quantification of the number of *Slc1a2^+^* astrocytes in a large region of interest using Imaris. Each point = 1 mouse (n=4-5 mixed male and female mice per genotype). Unpaired two-tailed student’s t-test on mouse average. **H)** Quantification of the number of *Mfge8* mRNA spots in the hippocampus shown as percent distribution on *Slc1a2^+^* astrocytic nuclei. Each point = mouse average (n=4-5 mixed male and female mice per genotype). Two-way ANOVA (Interaction: F (2, 21) = 6.188, p<0.01) followed by Bonferroni’s multiple comparisons post-hoc test on Y=log(Y) transformed mouse average. **I)** Representative images of MFG-E8 (white) immunoreactivity in the 6-mo hippocampal CA1 SR in WT and NL-F KI mice. Scale bar = 30 μm. **J)** Quantification of MFG-E8 immunoreactivity in large region using ImageJ. Each point = mouse average (n=7 male mice per genotype). Unpaired two-tailed student’s t-test on mouse average. **K)** Western blot probing for MFG-E8 (55-75 kDa) with respect to GAPDH loading control (39 kDa) in 6-mo WT and NL-F KI hippocampal homogenates. 1 lane is 1 mouse. **L)** Quantification of MFG-E8/GAPDH using ImageJ densitometry analysis. Each point = mouse average (n=5 male mice per genotype). Unpaired two-tailed student’s t-test on mouse average. **M)** Representative images of RNAScope followed by immunostaining probing for either *Itgav* (cyan) and *Itgb5* (magenta) integrin mRNA subunits (left) or *Itgav* (cyan) and *Itgb3* (magenta) integrin mRNA subunits (right) with microglial P2Y12 (green) in the hippocampus. Scale bar = 20 (left) and 15 (right) μm. Inset highlight ROIs. **N)** Flow cytometry probing for CD61 integrin on CD45^+^CD11b^+^CX3CR1^+^ hippocampal microglia from 6-mo WT and NL-F KI mice. **O)** Quantification of the levels of CD61 on CD45^+^CD11b^+^CX3CR1^+^ microglia. Each point = mouse average (n=5 female mice per genotype). Unpaired two-tailed student’s t-test followed by a Welch’s correction on mouse average. All data shown as mean ± SEM. p-values shown ns P>0.05; *P<0.05; **P<0.01; ***P<0.001; ****P<0.0001.

### Increased levels of the MFG-E8 integrin pathway when synapses are vulnerable to loss in the NL-F KI hippocampus

Given that we found significantly higher MFG-E8-immunoreactivity in the local milieu of bulbous astrocytes compared to bushy astrocytes in the NL-F KI hippocampus **(Figs 3J** and **K)**, we next asked whether astrocytic MFG-E8 is upregulated at the onset of synapse loss. Using RNAScope, we found increased *Mfge8* mRNA in astrocytes of 6-mo NL-F KI hippocampus compared to WT controls **(Figs 4E** and **F)**. We further assessed *Mfge8* mRNA expression using RT-qPCR from FACS-sorted cells, which showed an increasing trend of *Mfge8* in ACSA-2^+^CD45^-^ astrocytes from NL-F KI mice compared to WT controls with no difference in CD45^+^CD11b^+^CX3CR1^+^ microglia between the two genotypes **(Fig S6G)**. There was no change in the number of *Slc1a2*^+^ astrocytes between WT and NL-F KI mice **(Fig 4G)**, suggesting that the increase of *Mfge8* is driven by a change in expression rather than astrocyte number. Indeed, a greater proportion of *Slc1a2*^+^DAPI^+^ astrocytes expressed more *Mfge8* mRNA puncta in the NL-F KI hippocampus relative to WT controls **(Fig 4H)**. We confirmed that MFGE8 levels are also increased at the protein level in the NL-F KI hippocampus vs. WT controls by IHC **(Figs 4I** and **J)** and western-blotting **(Figs 4K** and **L)**.

MFG-E8 promotes engulfment by binding to *Itga_v_Itgb_3_* and *Itga_v_Itgb_5_*, integrin receptors on phagocytes^26^, which in the brain are expressed by P2Y12^+^ microglia **(Fig 4M)**^58^. In line, we observed higher levels of CD61 integrin on CD45^+^CD11b^+^CX3CR1^+^ microglia in the 6-mo NL-F KI hippocampus as compared to WT controls using flow cytometry **(Figs 4N** and **O)**. These results altogether indicate a region-specific increase of astrocytic MFG-E8 near bulbous astrocytes and an increase of the microglial integrin pathway, suggesting a potential for astrocyte-microglia crosstalk in modulating synaptic engulfment.

### MFG-E8 is necessary for Aβ oligomer-induced microglia-synapse engulfment and synapse loss *in vivo*

Next, we asked whether MFG-E8 is necessary for microglia-synapse engulfment *in vivo*. To accomplish this, we utilized a well-established *in vivo* model of Aβ oligomer-induced synapse loss in WT and *Mfge8* KO mice, where we administer synthetic humanized amyloid-β 40-S26C oligomers versus PBS control using intracerebroventricular (ICV) injection ^7,9,65–68^. At 18 and 72 hours (hr) post ICV injection, we assessed levels of microglia-engulfed synaptic material using high-resolution confocal imaging for Imaris 3D surface rendering and numbers of excitatory synapses using super-resolution Airyscan confocal microscopy, respectively, in the contralateral hippocampus **(Fig S7A)**. In line with published studies, we found that Aβ oligomers induced a significant increase of internalized Homer1 inside CD68^+^ lysosomes of P2Y12^+^ microglia in the contralateral CA1 SR hippocampus of WT mice vs. PBS controls **(Figs S7B-C)**; however, we did not see a similar Aβ oligomer-induced effect on engulfed synapse levels in microglia of *Mfge8* KO mice. To assess excitatory synapse loss, we quantified the colocalization between pre- and post-synaptic markers Bassoon and Homer1, respectively, in the hippocampal CA1 SR using super-resolution Airyscan confocal microscopy **(Fig S7D)**. While Aβ oligomers induced a significant loss of colocalized excitatory synaptic puncta in WT mice as compared to PBS controls, there was no significant difference in synapse numbers between Aβ injected vs. control-injected *Mfge8* KO mice **(Figs S7D-E)**. We validated that there were no appreciable changes in the number of NeuN^+^ neurons, Iba1^+^ microglia or GFAP^+^ astrocytes in *Mfge8* KO mice **(Fig S8)**. These results suggest that MFGE8 is required for Aβ oligomers to promote microglial engulfment of excitatory post-synaptic Homer1 and to induce synapse loss.

### CRISPR-saCas9 deletion of astrocytic MFG-E8 rescues microglia-synapse engulfment and loss in two genetic amyloidosis mouse models

Our results altogether suggest that astrocytes are the major cellular expressors of MFG-E8 in the brain parenchyma **(Fig 4)** and that MFG-E8 promotes microglia-synapse engulfment *in vitro* and *in vivo* **(Figs S4; S7)**. To confirm the role of astrocytic MFG-E8 in modulating microglia-synapse engulfment and synapse loss *in vivo*, we generated a viral CRISPR-saCas9 deletion strategy to specifically knock-down *Mfge8* from astrocytes **(Fig S9)**^34^. saCas9-HA and *Mfge8*- and *Ctrl-*gRNA-mCherry plasmids were packaged into AAV5 viral constructs under the astrocytic *GfaABC_1_D* promoter **(Fig S9)** (see methods). First, we evaluated whether the CRISPR-saCas9 paradigm effectively targets astrocytes and induces a knock-down of MFG-E8. Mice were unilaterally injected in the hippocampus with either control CRISPR (saCas9-HA alone or saCas9-HA with *Ctrl*-gRNA-mCherry) or test CRISPR (Cas9-HA and *Mfge8*-gRNA-mCherry) constructs **(Figs 5A; S10A)**. We assessed the colocalization of both *Ctrl*- and *Mfge8*-gRNA-mCherry constructs (mCherry) with cell-specific markers in the hippocampus **(Fig 5B)** and observed >90% of the mCherry-labeled signal colocalized with GFAP^+^ astrocytes with little to no colocalization with Iba1^+^ microglia or NeuN^+^ neurons (**Fig 5C)**. Furthermore, we found that >90% of GFAP^+^ astrocytes colocalized with mCherry-labeled signal upon injection with both control and test CRISPR (**Fig 5D)**, suggesting that the constructs target a significant proportion of hippocampal astrocytes with a high degree of specificity. In line with this finding, we found a >90% decrease in MFG-E8 protein levels by IHC in the mCherry-labeled zone in the hemispheres injected with test CRISPR **(Fig 5A)**, in comparison to non-injected or control CRISPR injected hemispheres across all mice tested **(Figs 5E** and **F)**, suggesting that the CRISPR-saCas9 paradigm successfully results in a robust and consistent knock-down of MFG-E8.

**Fig 5.**
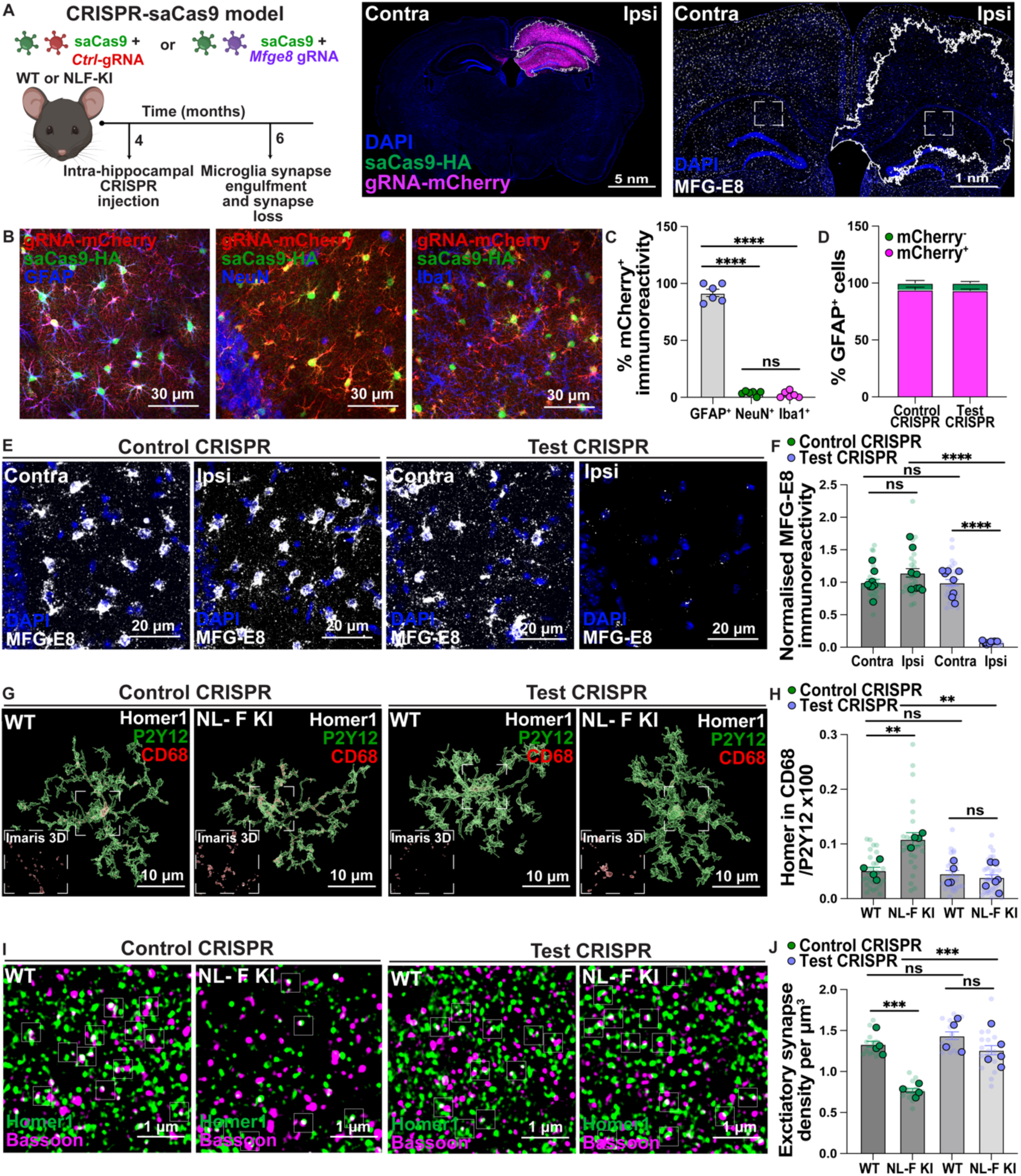
Viral CRISPR-saCas9 deletion of astrocytic MFG-E8 prevents microglia-Homer1 engulfment and excitatory loss in the NL-F KI hippocampus. **A)** Schematic of CRISPR-saCas9 paradigm to knock-down *Mfge8* from hippocampal astrocytes with spatio-temporal control. Immunostaining for gRNA-mCherry (magenta) and Cas9-HA (green) constructs post-CRISPR-saCas9 injection. Scale bar = 5 nm. Representative image of MFG-E8 (white) immunoreactivity post-CRISPR-saCas9 injection. Scale bar = 1 nm. Inset highlights ROIs. mCherry injection zone drawn with white ROI on ImageJ. **B)** Representative images of gRNA-mCherry (red) and Cas9-HA (green) constructs with either GFAP (blue, left), NeuN (blue, middle) or Iba1 (blue, right) post-CRISPR-saCas9 injection. Scale bar = 30 μm. **C)** Quantification of the colocalized mCherry-labeled signal with GFAP^+^ astrocytes, NeuN^+^ neurons or Iba1^+^ microglia in the hippocampus using ImageJ shown as percentage of the total mCherry-labeled signal. Each point = 1 mouse (n=6 mixed male and female mice). Brown-Forsythe (643.8 (2.000, 6.920, p<0.0001) and Welch ANOVA (339.9 (2.000, 8.829), p<0.0001) with Dunnett’s T3 multiple comparisons test. **D)** Quantification of the colocalized GFAP^+^ astrocytes with mCherry-labelled signal in the hippocampus using ImageJ shown as percentage of the total GFAP^+^ astrocytes. 8-10 male mice sampled. **E)** Representative images of MFG-E8 (white) immunoreactivity in the 6-mo WT and NL-F KI SR hippocampus post control or test CRISPR-saCas9 injection. Scale bar = 20 μm. **F)** Quantification of MFG-E8 immunoreactivity using ImageJ. Data shown as normalized to contra control CRISPR injected hemisphere. Transparent points = individual ROIs (3 ROIs per condition; 24 ROIs sampled in total per condition), full points = mouse average of ROIs (n=8 male mice per condition). Two-way ANOVA (Interaction: F (1, 28) = 218.6, p<0.0001) followed by Bonferroni’s multiple comparisons post-hoc test on Y=log(Y) transformed mouse average. **G)** Representative 3D rendered images of excitatory post-synaptic Homer1 (white), CD68 lysosomes (red) and P2Y12 (green) in the 6-mo WT and NL-F KI CA1 SR hippocampus post control or test CRISPR-saCas9 injection. Scale bar = 10 μm. Inset shows representative zoom of Homer1 inside CD68^+^ microglial lysosomes. **H)** Quantification of microglial Homer1 engulfment using Imaris 3D surface rendering shown as: Homer1 volume in CD68^+^ lysosomes in P2Y12^+^ microglial surface/P2Y12 volume x100. Transparent points = individual microglia (5-6 microglia per mouse; 24 WT control, 24 NL-F KI control, 24 WT test and 34 NL-F KI test injected microglia were sampled in total), full points = mouse average of ROIs (n=4-6 male mice per condition). Two-way ANOVA (Interaction: F (1, 14) = 4.929, p<0.05) followed by Bonferroni’s multiple comparisons post-hoc test on Y=log(Y) transformed mouse average. **I)** Representative images of excitatory post-synaptic Homer1 (green) and pre-synaptic Bassoon (magenta) immunostaining in the 6-mo WT and NL-F KI CA1 SR hippocampus post control or test CRISPR-saCas9 injection using super-resolution Airyscan confocal microscopy. Scale bar = 1 μm. Insets show regions of colocalization. **J)** Quantification of the number of colocalized Homer1 and Bassoon spots shown as density per µm^3^ using Imaris. Transparent points = individual ROIs (3 ROIs per mouse; 12 WT control, 12 NL-F KI control, 12 WT test and 15 NL-F KI test injected ROIs were sampled in total), full points = mouse average of ROIs (n=4-5 male mice per condition). Two-way ANOVA (Interaction: F (1, 13) = 11.02, p<0.01) followed by Bonferroni’s multiple comparisons post-hoc test on Y=log(Y) transformed mouse average. All data shown as mean ± SEM. p-values shown ns P>0.05; *P<0.05; **P<0.01; ***P<0.001; ****P<0.0001.

Next, we aimed to test the functional role of astrocytic MFG-E8 in synapse loss in the NL-F KI genetic amyloidosis mouse model^25^. Mice were injected at 4 mo, 8 weeks before the earliest reported synapse loss^8,9^ to allow for sufficient recovery time, and collected at 6-mo. In control CRISPR injected mice, there was a significant increase in the internalized Homer1 in CD68^+^ lysosomes of P2Y12^+^ microglia in NL-F KI CA1 SR hippocampus compared to WT mice, as well as a decrease in colocalized excitatory synaptic puncta **(Figs 5G-J)** in line with previous findings^8,9^. Conversely, we found near complete amelioration to WT levels in both the amounts of microglia-engulfed synapses as well as excitatory synapse numbers in the test CRISPR-injected NL-F KI mice **(Figs 5G-J)**. In parallel, we performed the same experiments in a more aggressive transgenic model, the hAPP J20 (J20) transgenic (Tg)^69^, where synapse loss is observed at 3 mo^7^. Here we injected the test or control constructs at 1.5 mo. Akin to the NL-F KI mice, we observed similar rescue of microglia Homer1 engulfment and in the number of excitatory synaptic puncta in test CRISPR injected J20 Tg mice as compared to control CRISPR or non-injected mice **(Fig S10).** Altogether, these data demonstrate that knockdown of MFG-E8 specifically from astrocytes prevents Aβ-induced microglia-Homer1 engulfment as well as excitatory synapse loss in pre-plaque stages in two chronic genetic mouse models of amyloidosis, suggesting a critical role for astrocytes and MFG-E8 in microglia-mediated synapse elimination in the earliest stages of amyloidosis.

## Discussion

A fundamental question in neurobiology is what mediates synapse loss in neurologic diseases such as AD^1–6^. Multiple studies have shown that microglia mediate synapse loss by engulfing synapses^7–16^; however, the precise cellular and molecular mechanisms remain elusive. Here, we propose that astrocytes act as cellular and molecular triggers of microglia-synapse elimination in pre-plaque stages of amyloidosis. We report an early and highly region-specific alteration of a subset of hippocampal astrocytes, specifically in their peri-synaptic processes, at onset of synapse loss. We found that the local milieu of these astrocytic peri-synaptic process has increased levels of the phagocytic modulator MFG-E8^26^ accompanied by increased microglia synapse engulfment and synapse loss. Using various *in vitro* and *in vivo* approaches, including a viral CRISPR-saCas9 system, we show that MFG-E8, secreted by astrocytes, is both sufficient and necessary for microglia to engulf synapses and that knockdown of astrocytic MFGE8 prevents synapse loss in multiple models of Aβ-induced synapse pathology. Our findings altogether propose a novel glia-immune-synapse interaction in the earliest stages of amyloidosis and nominate astrocytes and MFG-E8 as mediators of microglia-synapse elimination.

The study of astrocytes has relied on GFAP immunostaining, which only labels ∼15% of the astrocyte territory^32,35,70,71^. This conventional approach has led to the characterization of morphological and functional astrocyte changes in disease broadly termed ‘GFAP^+^ reactive gliosis’. Using more intricate tools, such as viral tdTomato labeling of astrocytes^33^ and super-resolution microscopy^48,49^, we found a distinct subset of astrocytes in the NL-F KI hippocampus with bulbous processes, which have a reduced territory and decreased synaptic coverage. We found that this subset begins to accumulate at the onset of synapse loss in the 6-mo NL-F KI hippocampus^8,9^ and that there is an exacerbated loss of excitatory post-synaptic Homer1 near the vicinity of bulbous processes. Previous studies have reported alterations in astrocyte morphology and astrocyte-synapse associations with age and in mouse models of neurodegenerative disease, such in the R6/2 Huntington’s disease model, APP/PS1 Tg amyloidosis AD model and SOD1 amyotrophic lateral sclerosis model as well as in human post-mortem and surgically resected tissue^34,42,48,72–74^. These emerging data propose changes in astrocyte morphology and synapse coverage as a potential shared mechanism of ageing and neurodegenerative disease, raising the critical question of whether diminished astrocytic support could act as a trigger for region-specific synapse loss.

Interestingly, we found that the bulbous astrocytes predominantly accumulate in the hippocampus^38,42^, suggesting that region-specific heterogeneity, either in steady-state or disease-associated, may underlie this phenotype. While we found an increase in the subset of bulbous astrocytes with progressive amyloidosis, interestingly, we did not observe a direct spatial association between bulbous astrocytes and Aβ plaques, suggesting that this astrocyte subset is not peri-plaque^43^ and likely not directly exacerbated by amyloidosis. Future experiments are needed to delve into upstream mechanisms inducing these bulbous astrocytes as well as cell intrinsic vs extrinsic mechanisms. A possible mechanism could be Aβ dysregulation of neuronal activity, which is an early hallmark of AD^65–67,75–79^. Astrocytes are poised to sense neuronal cues and could therefore respond to synaptic malfunction^80^. In turn, the reduced territory of the astrocytes accompanied by the bulbous processes formation could result in glutamate spillover and excitotoxicity, creating a dysfunctional loop^53^. Alternatively, as astrocyte end-feet are critical components of the blood brain barrier (BBB), either CNS- or peripherally derived blood-borne factors, which can infiltrate because of BBB breakdown in AD^81–83^, may modulate this astrocyte phenotype. Since we also found this subset accumulating with normal ageing in the WT hippocampus, common mechanisms between age, which is the greatest risk factor for AD^84^, and AD pathology, may underly this astrocytic phenotype.

Further investigation into how this astrocyte subset differs could provide deeper insights into the triggers and consequences of these cells in AD pathogenesis. However, the specific association of these astrocytes with p62 suggests that they might be linked to autophagy and ubiquitin proteosome pathways, which have been implicated in AD pathogenesis^85–87^. Astrocytic p62-immunreactive bodies, akin to those observed in Lafora bodies, polyglucosan bodies and corpora amylacea with age and disease have previously been reported^36–42^; however, their functional implications remain poorly understood. Here, we report increased levels of synapse loss near the vicinity of these bulbous astrocytes, concomitant with elevated levels of the phagocytic modulator MFG-E8. Interestingly, while we observed increased microglia-synapse engulfment in the vicinity of bulbous astrocytes, there were no changes in the number of microglia in this local milieu, unlike what has been shown for Aβ plaque-enriched stages and injury^88^. This suggests that distinct microglial responses may underlie these phenotypes. Further cell type-specific delineation within unique cellular microenvironments will give insight into the underlying molecular mechanisms and functional consequences.

Previous reports have shown that astrocytes engulf synapses in development, adulthood and in AD mouse models (APP/PS1 Tg and TauP301S) via phagocytic pathways such as Megf10/Mertk^10,14,22,23,89^; however, we did not observe any changes between WT and NL-F KI mice, at least in the engulfment of excitatory post-synaptic maker Homer1 at pre-plaque timepoints. Conversely, we found that Homer1 engulfment at these early stages of amyloidosis is attributed to microglia, which have previously been reported to engulf synapses in various contexts in health and disease^7–16,90^. It could be that astrocytes engulf synapses that are not Homer1^+^, or that at later Aβ plaque stages when microglia are overburdened^91^ that astrocytes play a part, akin to what has been shown for apoptotic cells^92,93^. Instead, here we show, that in pre-plaque stages astrocytes induce microglia to engulf synapses in their local milieu via MFG-E8. We report that astrocytes and microglia upregulate MFG-E8 and integrin receptors, respectively, at time points coinciding with synapse loss in the NL-F KI hippocampus. In line, previous studies have reported a dysregulation of MFG-E8 levels in mouse models of AD pathology and AD patient tissue^59–62^. MFG-E8 has previously been shown to modulate phagocytosis by bridging the interaction between the canonical ‘eat-me’ signal phosphatidylserine (PtdSer) on apoptotic cells and the integrin receptor^14,26–28,61,63^. Furthermore, synapses can locally externalize PtdSer in health and disease^8,94–98^. These data altogether suggest that the removal of synapses and apoptotic cells may rely on shared mechanisms, depending on the context^99,100^. Further studies are needed to delineate role in PtdSer in astrocyte-microglia crosstalk and whether similar crosstalk mechanisms determine synapse vulnerability in development and other disease context.

Finally, we report an intriguing astrocyte-microglia crosstalk via MFG-E8 in regulating synapse numbers in amyloidosis brains and show a novel functional role for hippocampal astrocytic MFG-E8 *in vivo*. Comparable mechanisms of bidirectional astrocyte-microglia crosstalk have previously been demonstrated in developmental synaptic pruning via IL-33^24^ as well as in disease-induced states via C1q, TNFα and Il-1α^101^. Previous studies have shown that microglia engulf synapses in mouse models of AD pathology via the classical complement cascade (C1q/C3)^7,10,15^, osteopontin (SPP1)^9^ and triggering receptor expressed on myeloid cells 2 (TREM2)^8,102^. Given the intimate association between astrocytes and synapses, astrocytic MFG-E8 may act upstream of these processes, however, further work is necessary to elucidate whether and how astrocytic MFG-E8 functions within these pathways.

### Limitations of the study

Our study identifies astrocytes as key players in modulating microglia-mediated synapse elimination in pre-plaque stages of mouse models of amyloidosis, whereby changes in peri-synaptic astrocytic processes including increased levels of phagocytic modulator, MFG-E8, create local regions of increased synapse loss. Using astrocyte specific CRISPR-saCas9 deletion of *Mfge8*, we show for the first time a functional role of astrocytic MFG-E8 *in vivo* and demonstrate that astrocytic MFG-E8 is necessary for microglia-synapse engulfment. These data highlight a novel cellular mechanism in earliest stages of AD pathogenesis and identify MFG-E8 as a potential therapeutic candidate for combating synapse loss in AD. However, we acknowledge that our study has focused on synapse loss during earliest stages of amyloidosis mouse models (NL-F KI and J20 Tg); plaque-enriched stages may induce distinct gliosis states and non-glia mediated neurodegeneration. Further, these mice do not develop neurofibrillary tangles nor neurodegeneration^25,69^. There is no AD mouse model that fully recapitulates the phenotypes observed in AD patient brains. Further work is necessary to understand whether astrocytic MFG-E8 plays a role in tau-induced microglia-mediated synapse loss^10,15^. In addition, our study has not assessed whether the rescued synapses are functional. Thus, it will be critical to understand whether astrocytic MFG-E8 is ultimately a beneficial or detrimental processes at the circuit level using cognitive assays.

## Acknowledgements

We thank members of the Hong lab and UK DRI for critical input to the manuscript; Elena Ghirardello for managing mouse colonies; Dr. Nicholas Cade for microscopy; Dr. Martha Foiani for guidance with human staining, Dr. Mathieu Bourdenx for guidance with autophagy mechanisms and p62/SQSTM1 biology. Dr. Takashi Saito, Dr. Hiroki Sasaguri and Dr. Takaomi C. Saido (RIKEN Brain Science Institute) for provision of hAPP NL-F KI. Dr. Shigekazu Nagata (RIKEN Brain Science Institute) for provision of MFG-E8 KO cryopreserved sperm. We thank the donators for their brains and the Newcastle Brain Tissue Resource including Dr. Debbie Lett for provision of human tissue. Dr. Monica Kim at AstraZeneca UK for help with primary astrocyte cultures. We thank Dr. Annerieke Sierksma at VIB KU Leuven and VIB for statistical analysis advice. Schematics were created with BioRender.com.

## Author contributions

Conceptualization, DS., SH. Methodology, DS., LW., BSK., SH. Software, N/A. Validation, SAG., FP., TG., YZ., AK. Formal analysis, DS., SAG., FP., TG., SDS., YZ., AK. Investigation, DS., SAG., FP., TG., SDS., JRC., GC., YZ., AK. Resources, BSK., SH. Data curation, N/A. Writing-Original draft, DS., SH. Writing-Review and Editing, DS., SAG., FP., TG., SDS., LW., JRC., YZ., GC., AK., PM., OJF., BSK., SH. Visualization, DS., SH. Supervision, DS., OFJ., BSK., SH. Project administration, BSK., SH. Funding Acquisition, DS., BSK., OJF., SH.

## Figures and legends

**Fig S1.**
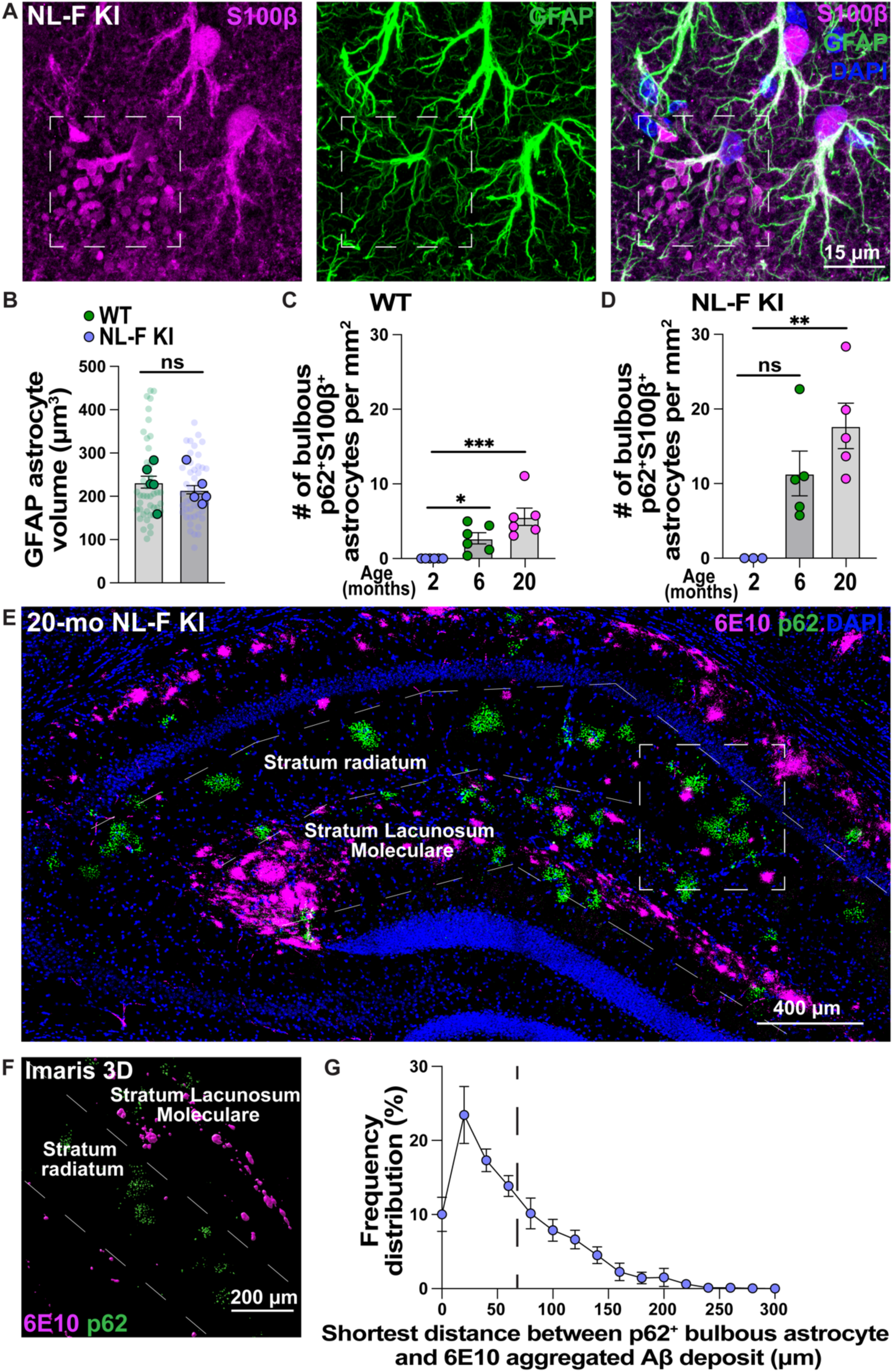
Bulbous astrocytes do not spatially correlate with Aβ plaques. **A)** Representative image of S100β (magenta) and GFAP (green) immunoreactivity in the hippocampus. Scale bar = 15 μm. Inset highlights exemplar of S100β^+^ processes that are not labeled by GFAP. **B)** Quantification of the GFAP^+^ astrocytic volume (μm^3^) using Imaris 3D surface rendering. Transparent points = individual astrocytes (7-11 astrocytes per mouse; 41 WT and 45 NL-F KI astrocytes sampled in total); full points = mouse average of ROIs (n=5 male mice per genotype). Unpaired two-tailed student’s t-test on mouse average. **C-D)** Quantification of the number of p62^+^ S100β^+^ bulbous astrocytes per mm^2^ in the WT **(C)** or NL-F KI **(D)** hippocampus. Each point = 1 mouse (n=3-6 male mice per age). Kruskal-Wallis (p<0.0001 **(C)** and p<0.01 **(D)**) test followed by the Dunn’s multiple comparisons test on mouse average. **E)** Representative image of p62 (green) and 6E10 (magenta) immunoreactivity in the 20-mo NL-F KI hippocampus. Scale bar = 400 μm. Dashed square represents ROI zoom for **F**. Dashes lines separated the stratum radiatum and stratum lacunosum moleculare. **F)** Imaris 3D reconstruction of p62- accumulated bulbous astrocytes (green) and 6E10-immunoreactive aggregated Aβ deposits (magenta). Scale bar = 200 μm. **G)** Quantification of shortest distance (μm) between p62-accumulated bulbous astrocytes and 6E10-labeled aggregated Aβ deposits. Data shown as frequency distribution (%) of p62- accumulated bulbous astrocytes (n=5 male mice). Dashed line represents the median distance. All data shown as mean ± SEM. p-values shown ns P>0.05; *P<0.05; **P<0.01; ***P<0.001; ****P<0.0001.

**Fig S2.**
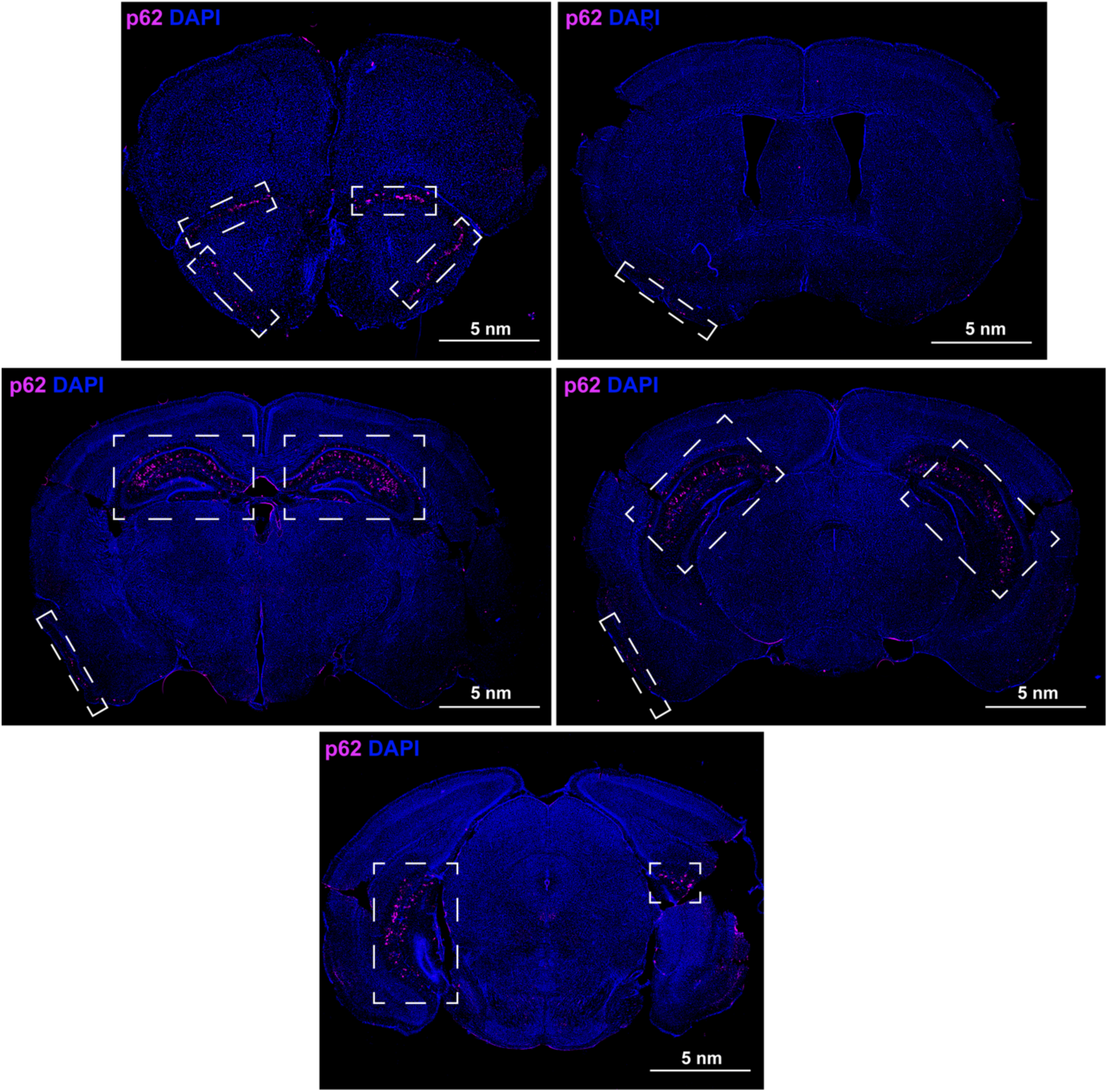
p62-accumulated bulbous astrocytes are found in a region-specific manner in the mouse cerebrum. **A)** Representative images of p62 (magenta) immunoreactivity in serial brain cross across the cerebrum in a 20-mo NL-F KI mouse. Only regions with bulbous p62-accumulated S100β^+^ astrocytes are represented i.e. piriform cortex, hippocampus and subiculum. Scale bar = 5 nm. Spatial patterning recapitulated in n=6 mixed age/genotype/sex mice. Insets highlight regions where bulbous astrocytes were found.

**Fig S3.**
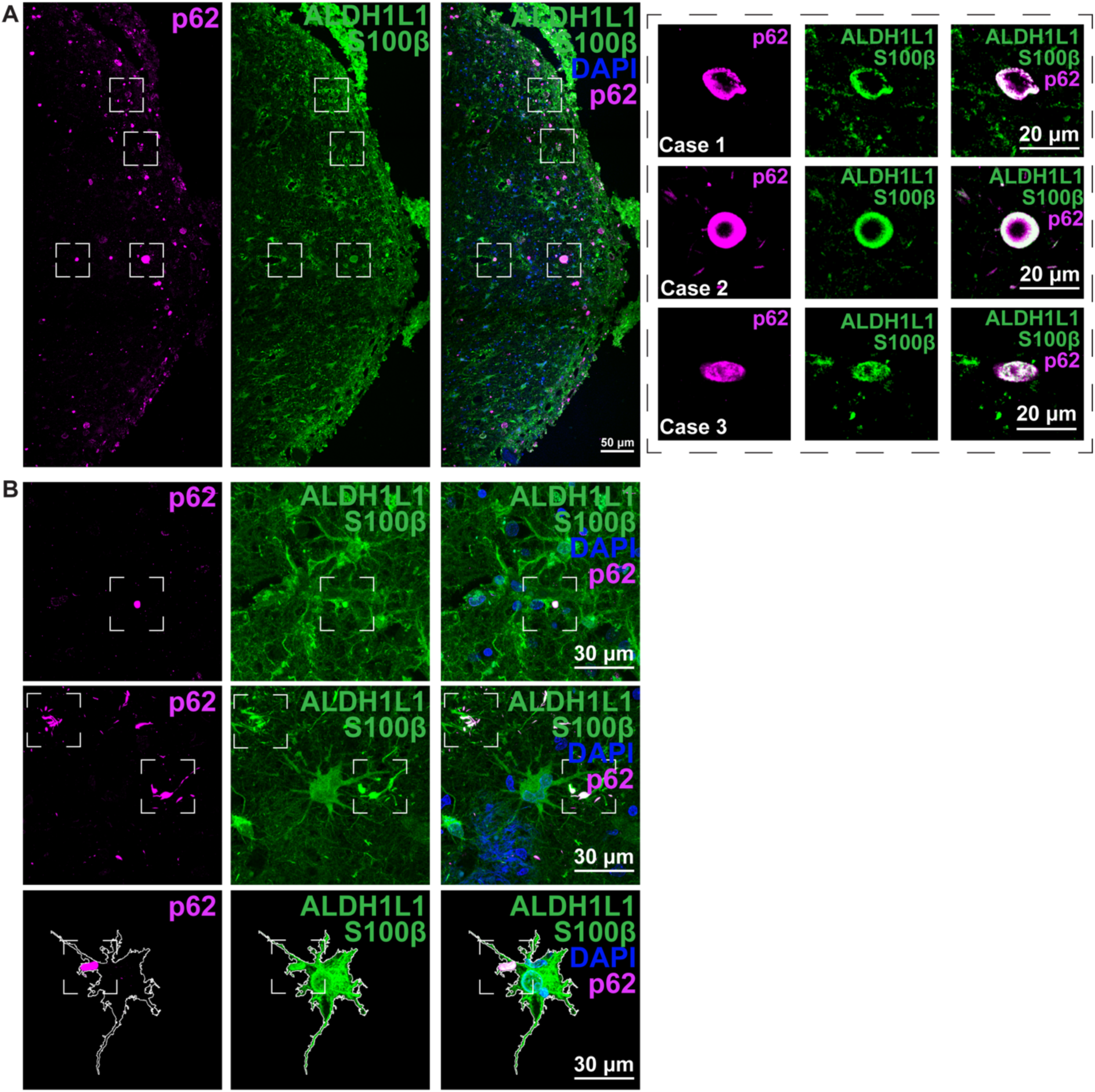
Analogous p62-accumulated astrocyte profile may exist in the human post-mortem hippocampus. **A)** Representative images of p62 (magenta) and S100β/ALDH1L1^+^ (green) astrocytes in the human hippocampus. Inset highlight representative zoom-ins of three different cases. Scale bar = 50, 20 μm. **B)** Representative images of p62 (magenta) and S100β/ALDH1L1^+^ (green) astrocytes in the human hippocampus. Scale bar = 30 μm. (n=2). Images are modified on ImageJ as shown by white ROI. Insets highlight ROIs. Representative images are shown from both control and AD patients. Please see **Table 1** for patient demographics.

**Fig S4.**
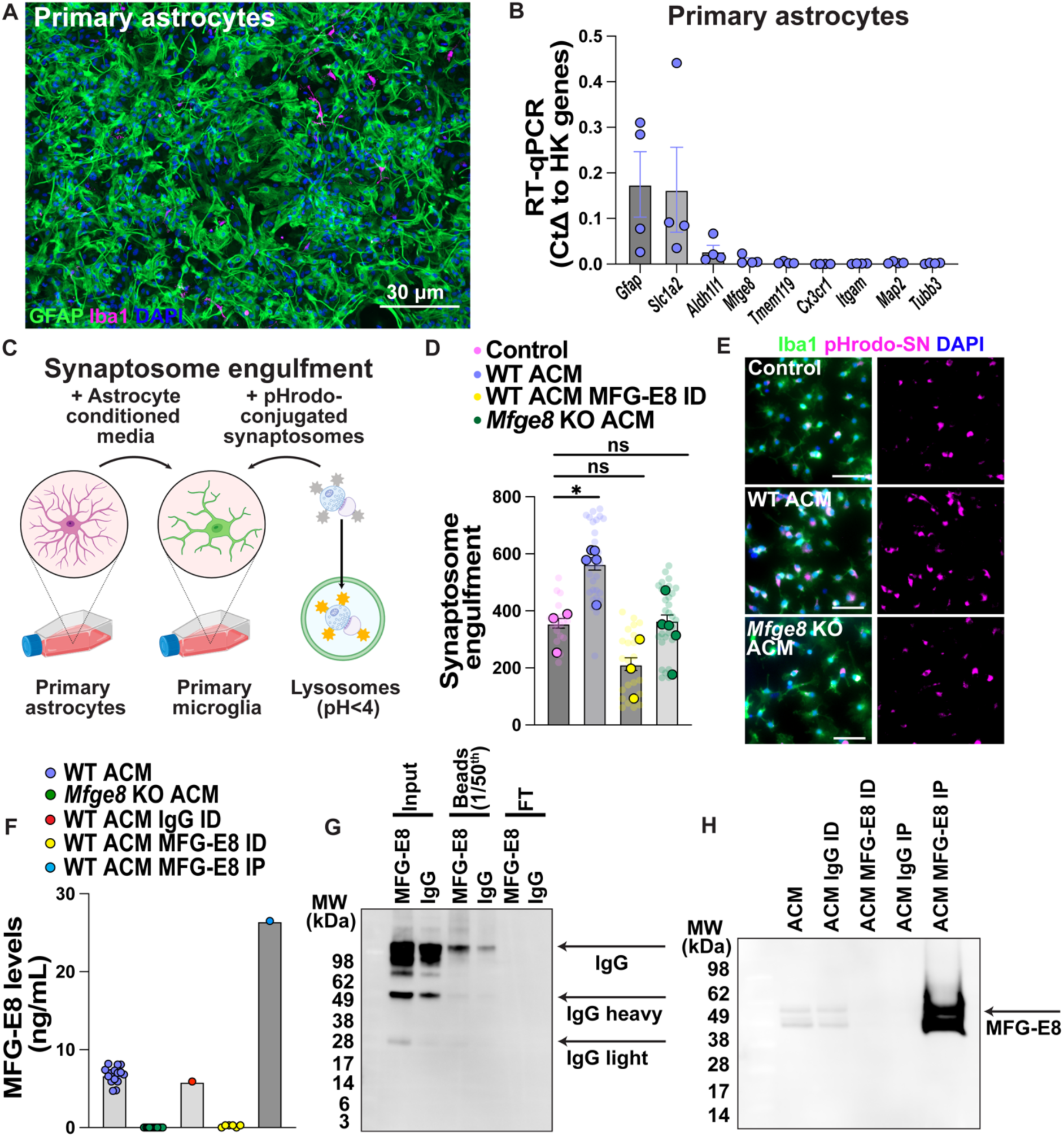
Secreted MFG-E8 in astrocyte conditioned media is sufficient to promote microglia- synaptosome engulfment. **A)** Representative image of GFAP (green) and Iba1 (magenta) immunoreactivity in primary astrocytic cultures. Scale bar = 30 μm. **B)** RT-qPCR probing for gene expression of *Gfap*, *Slc1a2*, *Aldh1l1*, *Mfge8*, *Tmem119*, *Cx3cr1, Itgam*, *Map2* and *Tubb3* in primary astrocytic cultures. Gene expression is normalized to the geomean of 3 housekeeping genes (*Actb*, *Gapdh* and *Rpl32*) by the Delta CT method. Each point = 1 well (n=4 wells). **C)** Schematic of primary microglial synaptosome engulfment assay using ACM collected from primary astrocytes. **D)** Quantification of the pHrodo fluorescence intensity with time shown as area under curve. Transparent points = individual ROIs with approx. 30 microglia per ROI (2-12 ROIs culture preparation; 17 control microglial media, 28 WT ACM, 29 *Mfge8* KO ACM and 22 WT ACM MFG-E8 ID ROIs sampled in total), full points = culture average of ROIs (n=3-5 mixed male and female culture preparations). One-way ANOVA followed (F (3, 12) = 10.86, p<0.001) by Bonferroni’s multiple comparisons post-hoc test on culture average. **E)** Representative images of Iba1 (green) immunoreactivity and pHrodo-synaptosomes (SN) (magenta) in primary microglial cultures treated with either control media, WT ACM or *Mfge8* KO ACM using widefield microscopy on the Celldiscoverer 7. Scale bar = 40 μm. **F)** Mouse MFG-E8 ELISA probing for levels of MFG-E8 (ng/mL) in WT ACM, *Mfge8* KO ACM and fractions from the MFG-E8 ID reaction (IgG ID, MFG-E8 ID and MFG-E8 IP). Each point = 1 culture well (15 WT and 15 MFG-E8 KO wells sampled in total across 2-3 independent culture preparations) or ID preparation (6 WT MFG-E8 ID ACM preparations sampled in total, 1 IgG ID and 1 MFG-E8 IP sampled as control). **G)** Western blot probing for goat IgG (150, 50 and 25 kDa) using a secondary anti-goat HRP in the ID reaction. The lanes represent the input antibody put into the reaction, 1/50^th^ of the total beads and the flow through (FT) for MFG-E8 and an IgG control reaction. **H)** Western blot probing for MFG-E8 (55 kDa) in the fractions from the ACM ID reaction (input, IgG ID, MFG-E8 ID, IgG IP and MFG-E8 IP). All data shown as mean ± SEM. p-values shown ns P>0.05; *P<0.05; **P<0.01; ***P<0.001; ****P<0.0001.

**Fig S5.**
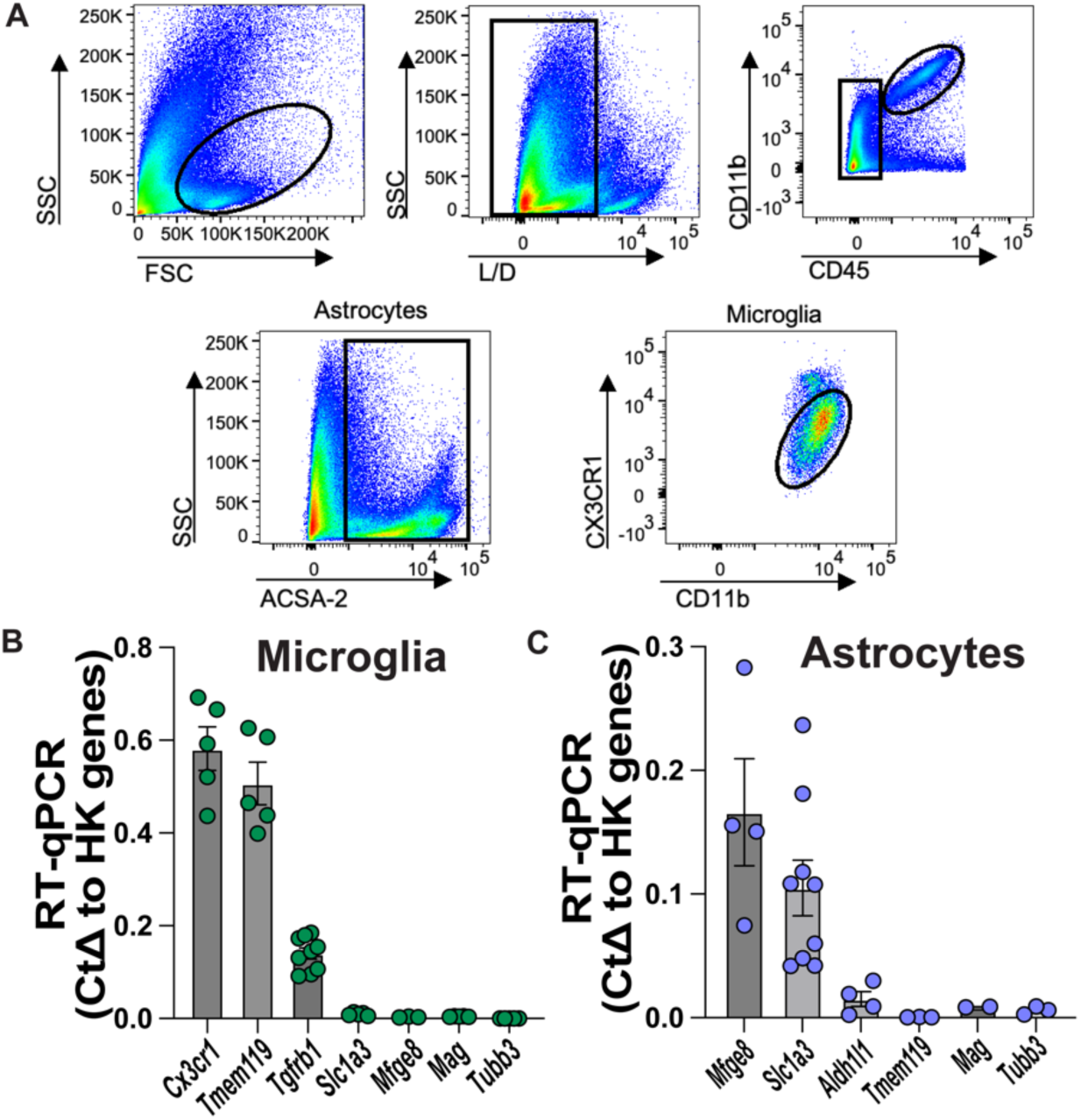
Validation and characterization of FACS cell sorting. **A)** Flow cytometry plots showing gating strategy based on forward (FSC), side scatter (SSC) and levels of ACSA-2, CD11b, CD45 and CX3CR1. Microglia are selected based on live and CX3CR1^+^CD45^+^CD11b^+^. Astrocytes are selected based on live and ACSA-2^+^CD45^-^. **B)** RT-qPCR probing for gene expression of *Cx3cr1*, *Tmem119*, *Tgfrb1*, *Slc1a3*, *Mfge8*, *Mag* and *Tubb3* on FACS sorted CX3CR1^+^CD45^+^CD11b^+^ microglia. Gene expression is normalized to the geomean of 3 housekeeping genes (*Actb*, *Gapdh* and *Rpl32*) by the Delta CT method. Each point = 1 mouse (n=3-9 female mice). **C)** RT-qPCR probing for gene expression levels of *Mfge8*, *Slc1a3*, *Aldh1l1*, *Tmem119*, *Mag* and *Tubb3* on FACS sorted ACSA-2^+^CD45^-^ astrocytes. Gene expression is normalized to the geomean of 3 housekeeping genes (*Actb*, *Gapdh* and *Rpl32*) by the Delta CT method. Each point = 1 mouse (n=2-9 female mice). All data shown as mean ± SEM.

**Fig S6.**
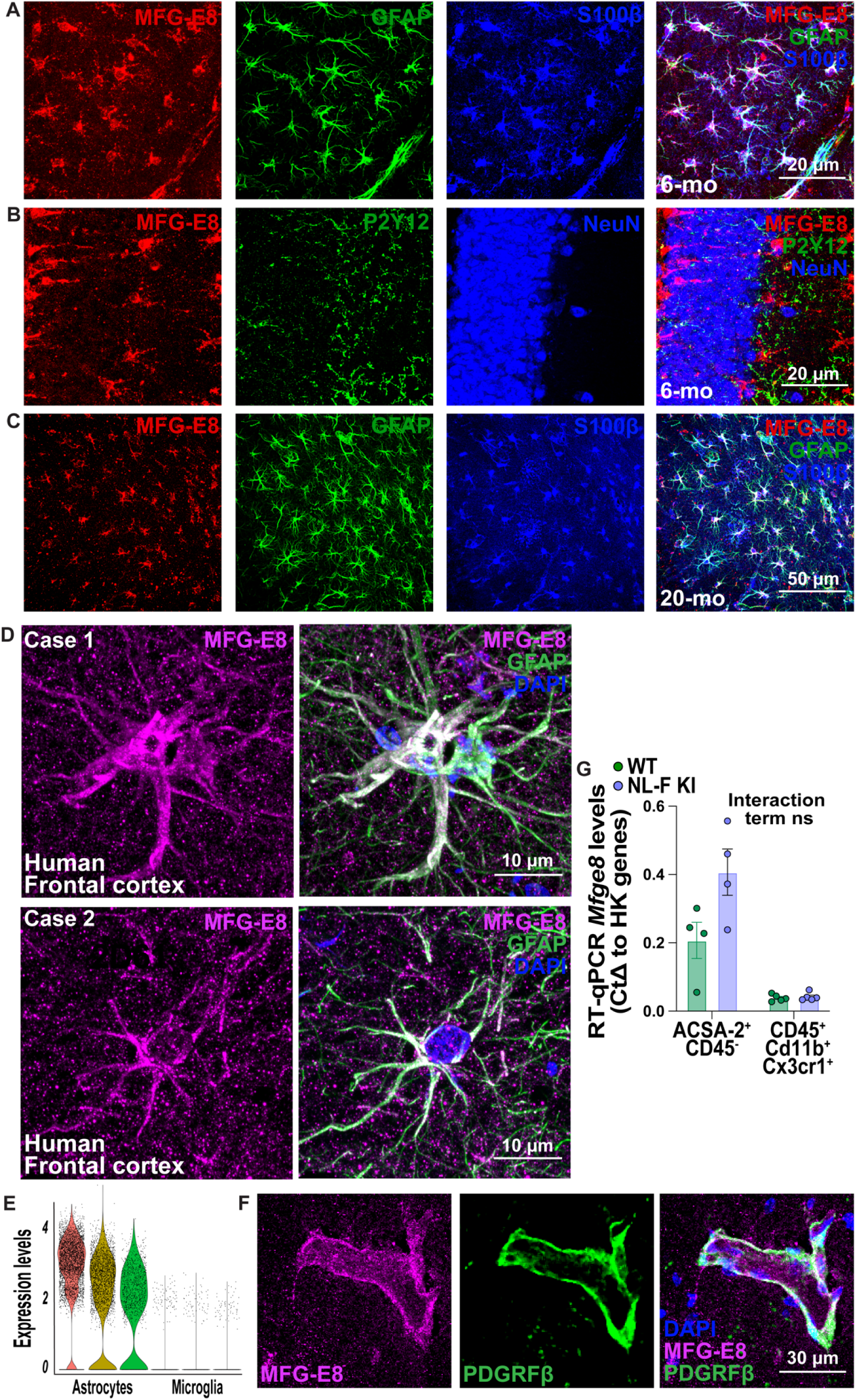
MFG-E8 is expressed by astrocytes in the mouse and human brain parenchyma. **A)** Representative image of MFG-E8 (red) and GFAP^+^ (green) and S100β^+^ (blue) astrocytes in the 6-mo mouse hippocampus. Scale bar = 20 μm. **B)** Representative image of MFG-E8 (red), P2Y12^+^ microglia (green) and NeuN^+^ (blue) neurons in the 6-mo mouse hippocampus. Scale bar = 20 μm. **C)** Representative image of MFG-E8 (red) and GFAP^+^ (green) and S100β^+^ (blue) astrocytes 20-mo mouse hippocampus. Scale bar = 20 μm. **D)** Representative image of MFG-E8 (magenta) and GFAP (green) immunoreactivity in the post-mortem human frontal cortex. Representative images are from control and AD patients. Please see **Table 1** for patient demographics. Scale bar = 10 μm. **E)** Single cell RNA sequencing data from the Linnarson database^55^ comparing *Mfge8* expression levels on astrocyte and microglial clusters. **F)** Representative image of MFG-E8 (magenta) and PDGFRβ^+^ (green) immunoreactivity in the mouse hippocampus. Scale bar = 30 μm. **G)** RT-qPCR probing for *Mfge8* gene expression in FACS sorted ACSA2^+^CD45^-^ astrocytes and CX3CR1^+^CD45^+^CD11b^+^ microglia from 6-mo WT and NL-F KI mice. Gene expression is normalized to the geomean of 3 housekeeping genes (*Actb*, *Gapdh* and *Rpl32*) by the Delta CT method. Each point = 1 mouse (n=4-5 female mice per genotype). Two-way ANOVA (Interaction: F (1, 14) = 2.956, ns p>0.05) on Y=log(Y) transformed mouse average. All data shown as mean ± SEM. p-values shown ns P>0.05; *P<0.05; **P<0.01; ***P<0.001; ****P<0.0001.

**Fig S7.**
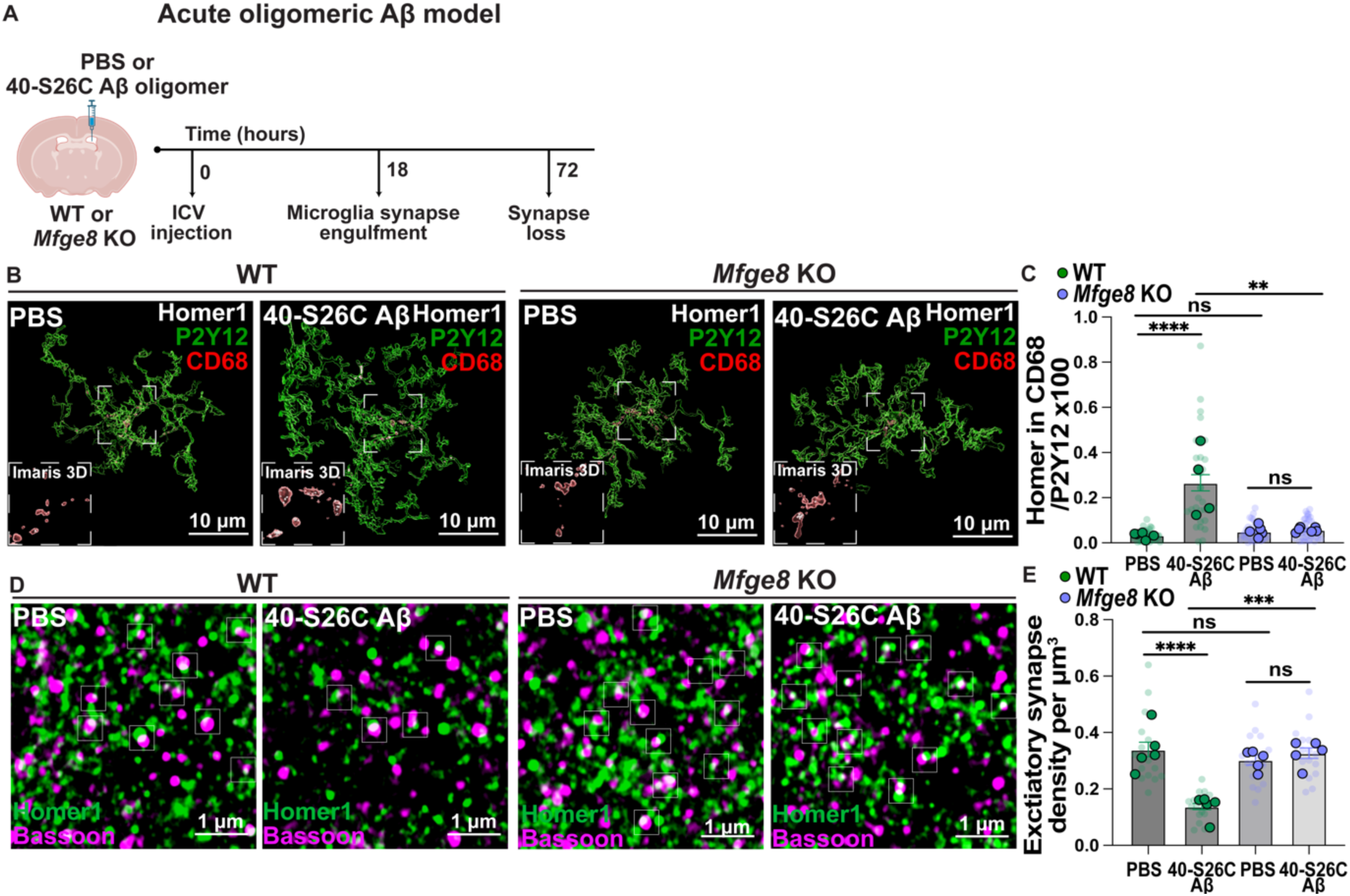
MFG-E8 is necessary for microglia-Homer1 engulfment and excitatory synapse loss in an *in vivo* acute Aβ injection model. **A)** Schematic of acute 40-S26C oligomeric Aβ ICV injection model. **B)** Representative 3D images for excitatory post-synaptic Homer1 (white), CD68 lysosomes (red) and P2Y12 (green) in the WT or *Mfge8* KO CA1 SR hippocampus post PBS control or synthetic humanized Aβ 40-S26C oligomer ICV injection. Scale bar = 10 μm. Inset shows representative zoom of Homer1 inside CD68^+^ microglial lysosomes. **C)** Quantification of microglial Homer1 engulfment using Imaris 3D surface rendering shown as: Homer1 volume in CD68^+^ lysosomes in P2Y12^+^ microglial surface/P2Y12 volume x100. Transparent points = individual microglia (5-9 microglia per mouse; 33 WT PBS, 28 WT Aβ, 31 *Mfge8* KO PBS and 44 *Mfge8* KO Aβ microglia were sampled in total), full points = mouse average of ROIs (n=4-6 female mice per condition). Two-way ANOVA (Interaction F (1, 16) = 17.39, p<0.001) followed by Bonferroni’s multiple comparisons post-hoc test on Y=log(Y) transformed mouse average. **D)** Representative images of excitatory post-synaptic Homer1 (green) and pre-synaptic Bassoon (magenta) immunoreactivity in the WT or *Mfge8* KO SR hippocampus post PBS control or synthetic humanized Aβ 40-S26C oligomer ICV injection using super-resolution Airyscan confocal microscopy. Scale bar = 1 μm. Insets show regions of colocalization. **E)** Quantification of the number of colocalized Homer1 and Bassoon spots shown as density per µm^3^ using Imaris. Transparent points = individual ROIs (3 ROIs per mouse; 15 ROIs were sampled per condition in total), full points = mouse average of ROIs (n=5 male mice per condition). Two-way ANOVA (Interaction: F (1, 16) = 23.29, p<0.001) followed by Bonferroni’s multiple comparisons post-hoc test on mouse average. All data shown as mean ± SEM. p-values shown ns P>0.05; *P<0.05; **P<0.01; ***P<0.001; ****P<0.0001.

**Fig S8.**
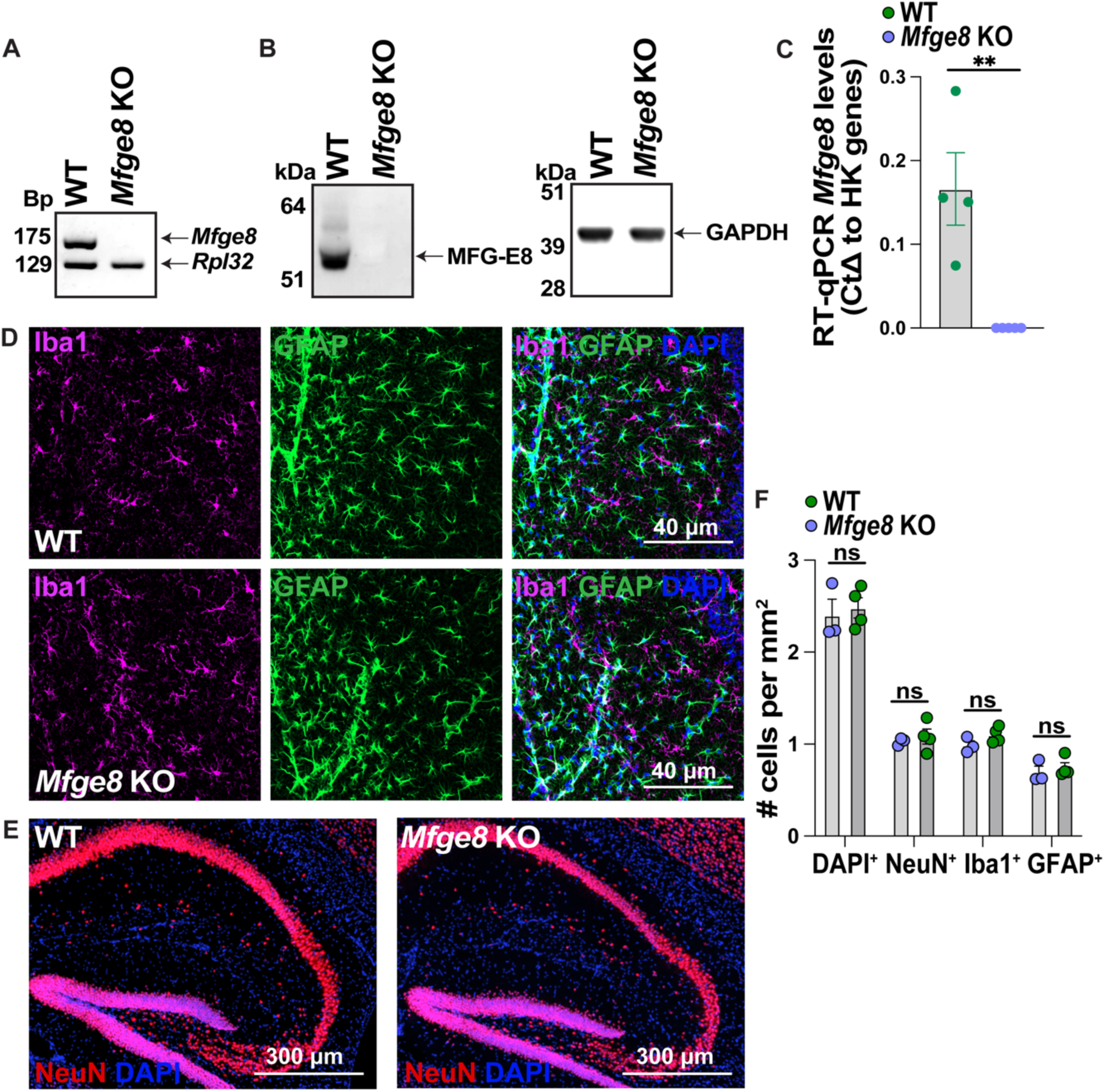
Validation and characterization of the *Mfge8* KO mouse model. **A)** PCR probing for *Mfge8* expression in hippocampal homogenate resolved on agarose gel with respect to housekeeping gene *Rpl32*. 1 lane = 1 mouse. **B)** Western blot probing for MFG-E8 (55-75 kDa) with respect to GAPDH loading control (39 kDa) in WT and *Mfge8* KO homogenate. 1 lane = 1 mouse. **C)** RT-qPCR probing for *Mfge8* gene expression in FACS sorted ACSA2^+^CD45^-^ astrocytes from WT and *Mfge8* KO mice. Gene expression is normalized to the geomean of 3 housekeeping genes (*Actb*, *Gapdh* and *Rpl32*) by the Delta CT method. Each point = 1 mouse (n=4-5 female mice per genotype). Mann-Whitney test on mouse average. **D)** Representative images of GFAP (green) and Iba1 (magenta) immunostaining in the hippocampus of WT and *Mfge8* KO mice. Scale bar = 40 μm. **E)** Representative images of NeuN (red) immunoreactivity in the hippocampus of WT and *Mfge8* KO mice. Scale bar = 300 μm. **F)** Quantification of the number of total DAPI^+^ cells, GFAP^+^ astrocytes, Iba1^+^ microglia and NeuN^+^ neurons in the hippocampus shown as number per mm^2^ using Imaris 3D surface rendering. Each point = 1 mouse (n=3-4 female mice per genotype). Multiple unpaired t-tests followed by Dunn’s multiple comparisons correction on mouse average. All data shown as mean ± SEM. p-values shown ns P>0.05; *P<0.05; **P<0.01; ***P<0.001; ****P<0.0001.

**Fig S9.**
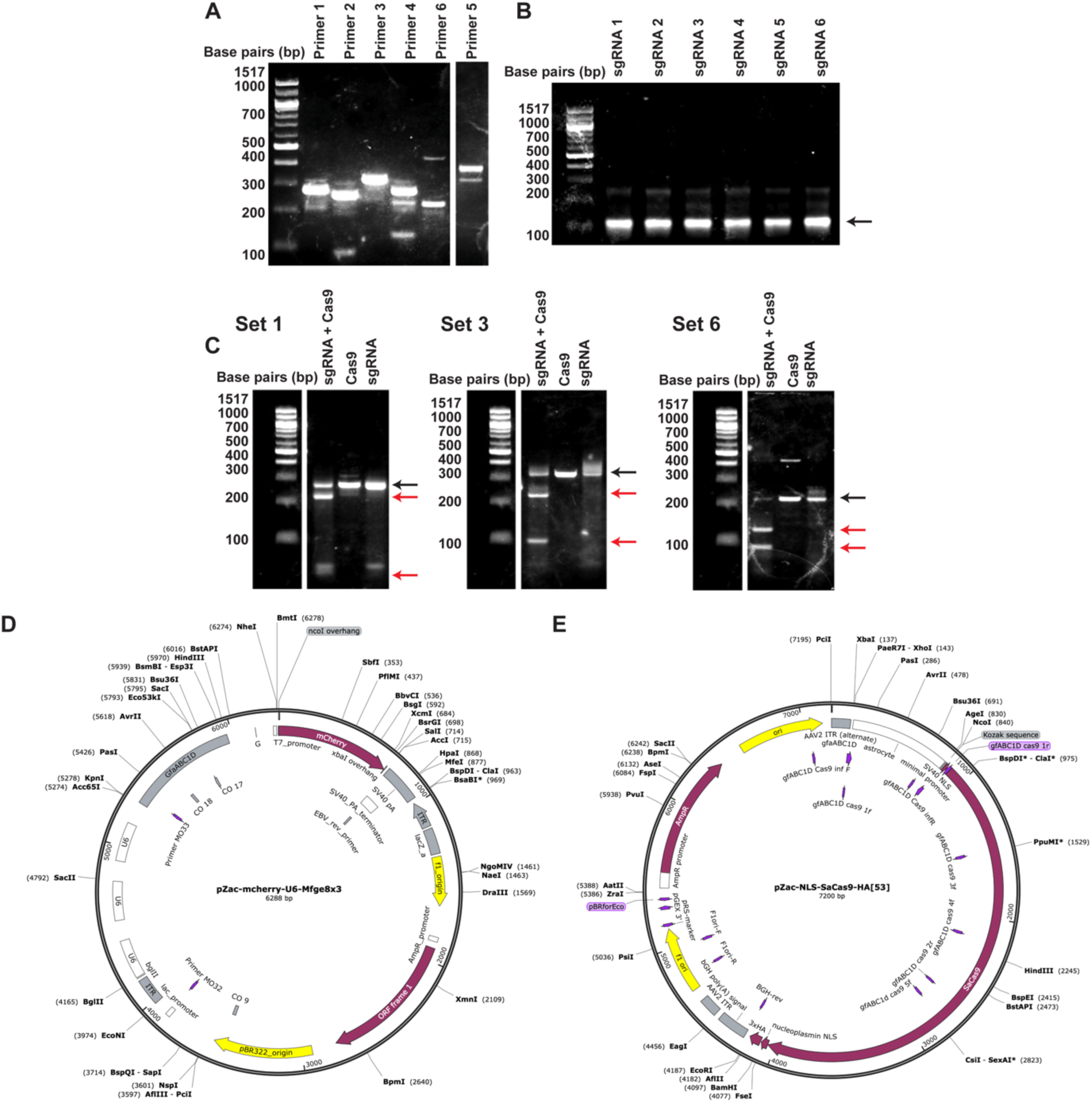
Design and testing of CRISPR-saCas9 paradigm. **A)** PCR product resolved on agarose gel after amplification of genomic DNA (gDNA) regions of the *Mfge8* gene. These gDNA amplicons correspond to the cleavage regions of the sgRNAs. Primer 1-6 refers to the primer set used to amplify each region. Expected amplicon product lengths: primer 1, 275 bp; primer 2, 254 bp; primer 3, 323 bp; primer 4, 275 bp; primer 5, 380 bp; primer 6, 232 bp. **B)** PCR product resolved on agarose gel after amplification of 6 DNA templates for subsequent transcription of the sgRNA molecules. Each template corresponds to one sgRNA sequence. The arrow points at the expected amplicon length (121 bp). **C)** *In vitro* digestion assay resolved on agarose gel to assess the cleavage efficiency of each designed sgRNA. The reaction was performed with the sgRNA + Cas9 or as a control the Cas9 or sgRNA alone. Black arrows indicate the input gDNA length and the red arrows indicate the expected cleavage fragments. Only the three successful reactions are shown here. Expected fragment lengths: Set 1, 218 bp and 31 bp; Set 3, 206 bp and 90 bp, Set 6, 127 bp and 104 bp. **D)** *Mfge8*gRNA-*GfaABC1D*-mCherry plasmid construct map. **E)** *GfaABC1D*-saCas9-HA plasmid construct map.

**Fig S10.**
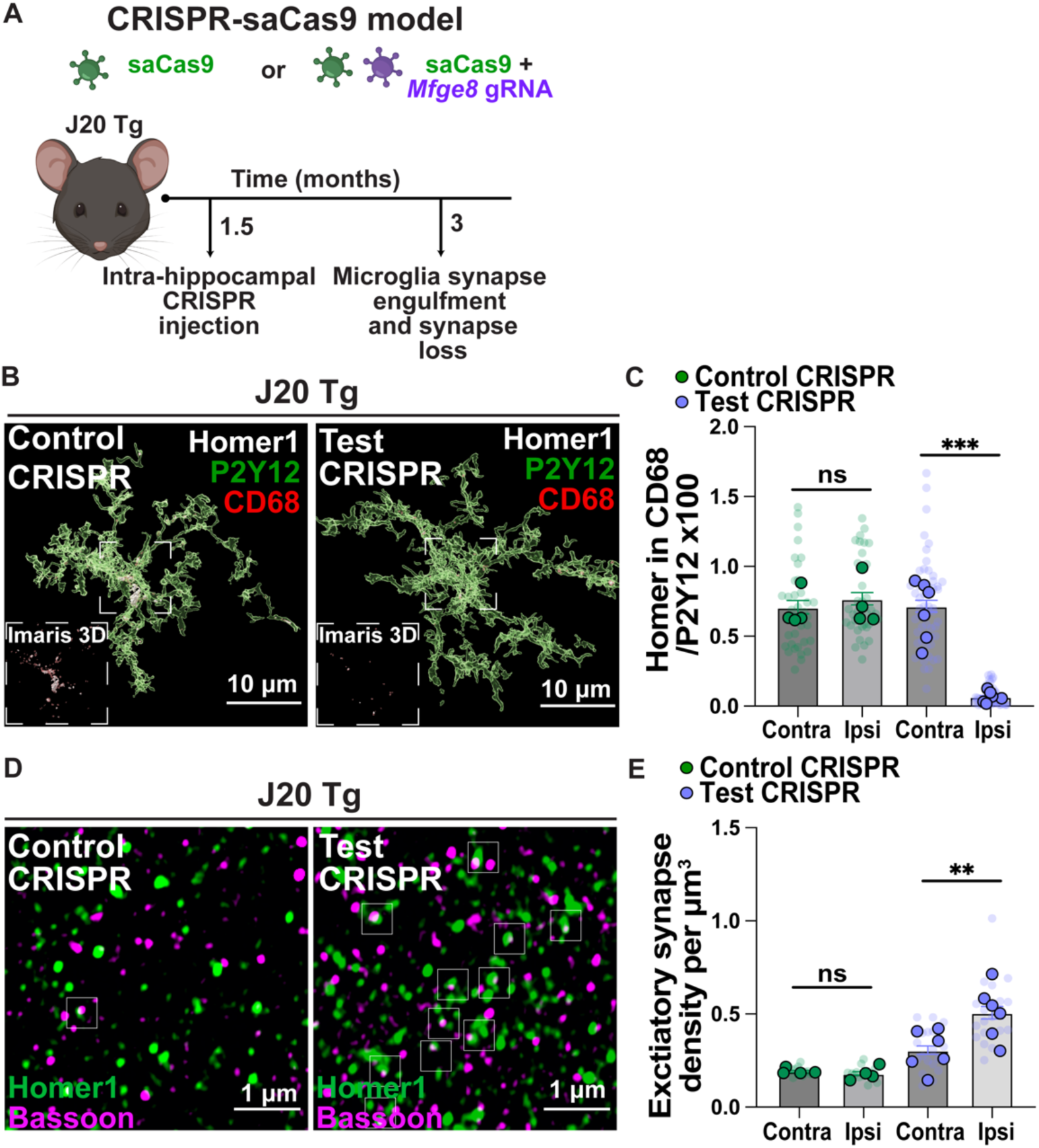
Preventative viral CRISPR-saCas9 deletion of astrocytic MFG-E8 rescues microglia- Homer1 engulfment and excitatory synapse loss in a second model of amyloidosis, the J20 Tg. **A)** Schematic of CRISPR-saCas paradigm to knock-down *Mfge8* from hippocampal astrocytes with spatio-temporal control. **B)** Representative 3D images for excitatory post-synaptic Homer1 (white), CD68 lysosomes (red) and P2Y12 (green) in the 3-mo J20 Tg CA1 SR hippocampus post control or test CRISPR-saCas9 injection. Scale bar = 10 μm. Inset shows representative zoom of Homer1 inside CD68^+^ microglial lysosomes. **C)** Quantification of microglial Homer1 engulfment using Imaris 3D surface rendering shown as: Homer1 volume in CD68^+^ lysosomes in P2Y12^+^ microglial surface/P2Y12 volume x100. Transparent points = individual microglia (6-11 microglia per condition; 34 contra control CRISPR, 34 ipsi control CRISPR, 51 contra test CRISPR and 46 ipsi test CRISPR injected microglia were sampled in total), full points = mouse average of ROIs (n=4-6 mixed male and female mice per condition). Multiple paired t-tests followed by the Bonferroni-Dunn correction on mouse average. **D)** Representative images of excitatory post-synaptic Homer1 (green) and pre-synaptic Bassoon (magenta) immunoreactivity in the 3-mo J20 Tg CA1 SR hippocampus post control or test CRISPR- saCas9 injection using super-resolution Airyscan confocal microscopy. Scale bar = 1 μm. Insets show regions of colocalization. **E)** Quantification of the number of colocalized Homer1 and Bassoon spots shown as density per µm^3^ using Imaris. Transparent points = individual ROIs (3 ROIs per condition; 12 contra control CRISPR, 12 ipsi control CRISPR, 18 contra test CRISPR and 18 ipsi test CRISPR injected ROIs were sampled in total), full points = mouse average of ROIs (n=4-6 mixed male and female mice per condition). Multiple paired t-tests followed by the Bonferroni-Dunn correction on mouse average. All data shown as mean ± SEM. p-values shown ns P>0.05; *P<0.05; **P<0.01; ***P<0.001; ****P<0.0001.

## Materials and Methods

### Animals

All experiments were performed in accordance with the UK Animal Scientific Procedures Acta 1986. Experimental procedures were approved by the UK Home Office and ethical approval was granted through consultation with veterinary staff at University College London (UCL). All animal work was completed under the appropriate ethics and licenses.

All animals were housed under temperature-controlled (temperature, 23.1 °C; humidity, 30– 60%) and pathogen-free conditions with 12 hr light/12 hr dark cycle with an ad libitum supply of food and water. Wild-type (WT) (C57BL/6J and C57Bl6/NCrl) mice were obtained from Charles River. hAPP NL-F KI (C57BL/6-App<tm2(NL-F)Tcs>) mice were originally provided by Takaomi Saido and distributed by Frances Edwards, UCL. hAPP J20 Tg (B6.Cg-Zbtb20Tg (PDGFB-APPSwInd)20Lms/2Mmjaxn) mice were provided by Lennart Mucke and distributed by Patricia Salinas, UCL. MFG-E8 KO (B6;129-Mfge8<TM1OSA>/OsaRbrc) were redrived onto the WT C57Bl6/NCrl background using imported sperm from RIKEN. Homozygous mice were used for all experiments.

Pilot experiments were conducted using both female and male mice to ensure there were no sex-specific phenotypes. Both male and female age-matched mice were then used throughout in this study to ensure there is no gender bias. For most of the experiments, a single sex was used at random depending on availability. For some experiments, both male and female mice were used. All are clearly stated where applicable in the figure legends. For example: male mice were used for spatiotemporal quantification of bulbous astrocytes, CRISPR-saCas9 NL-F KI studies and *Mfge8* KO 72 hr synapse loss studies. For example: female mice were used for: integrin flow cytometry, FACS sorting, *Mfge8* KO 18 hr microglia Homer1 engulfment studies. For example: both sexes were used for CRISPR-saCas9 J20 Tg studies and culture experiments. For all experiments, we ensured that we use appropriate matching controls re: genotype-, sex- and age.

For tissue harvest, mice were deeply anaesthetized with intraperitoneal pentobarbital. Mice were then transcardially perfused with either ice-cold PBS for biochemical assays or in combination with 4% PFA for immunostainings. For the latter, the brains were removed from the skull and post-fixed in 4% PFA for 24 hours at 4°C. After being rinsed in PBS, the brains were placed in 30% sucrose for up to 48 hours and frozen in tissue-embedding molds filled with OCT. Brain slices were sectioned either sagitally or coronally depending on the experiment using a cryostat microtome chamber (Leica CM1860 UV) at 12 μm thickness and mounted on glass slides (SuperFrost GOLD Adhesion Slides 11976299, Fisher Scientific) for RNAScope experiments or at 30 μm thickness and stored in cryoprotect solution at -20°C for free-floating immunohistochemistry.

### Primary microglial cultures

Primary microglial cultures were prepared as previously described^8,9^. Eight to ten pups of either sex aged P0-P4 per prep were decapitated. The brains were detached from the skull and the meninges were removed in ice-cold Hank’s Balanced Salt Solution (HBSS) with 1% penstrep (Gibco). The cortices and hippocampi were isolated and homogenized in HBSS using a P1000 pipette. The homogenate was put through a 70 μM pre-strainer and centrifuged at 400 g for 5 min at 4°C. The supernatant was removed, and the cell pellet was resuspended in ice-cold 35% isotonic percoll. The interface was carefully created with HBSS. The samples were centrifuged for 40 min at 4°C at 2,800 g with no break and with slow acceleration and deceleration. The myelin layer and supernatant layers were aspirated, and the cell pellet was washed in HBSS. Cells were counted and resuspended in microglial base media: DMEM F12 (Gibco), 5% fetal bovine serum (Gibco), 1% pen-strep. Cells were plated on borate buffer 0.1 M pH 8.5 poly-D lysine (Gibco) coated plates in 1 mL of microglial media at a density of 650k per well. Cells were maintained in an incubator at 37°C and 5% CO_2_. 90% of media was changed the day after and subsequently half of the media was changed every 2 days. After the initial 90% change, the media was supplemented with 50 ng/mL CSF1 (416-ML-010/CF RnD Systems), 50 ng/mL TGFβ1 (7666-MB-005/CF RnD Systems) and 100 ng/mL CX3CL1 472-FF-025/CF RnD Systems) to induce a more *in vivo*-like profile of microglia^103,104^.

### Primary astrocyte cultures

We established a primary GFAP^+^ astrocyte culture enriched in astrocytic transcripts (*Gfap*, *Aldh1l1* and *Slc1a3*) with little to no contamination from Iba1^+^ cells and microglial transcripts (*P2y12*, *Tmem119* and *Cx3cr1*) (**Fig S4)**. Eight to ten pups of either sex aged P0-P4 per prep were decapitated. The brains were detached from the skull and the meninges were removed in ice-cold Hank’s Balanced Salt Solution (HBSS) with 1% penstrep (Gibco). Once the cortices and hippocampi were isolated, they were digested in 0.25% Trypsin-EDTA (Gibco) and 200 Kunitz units/mL of DNAse I (ThermoFisher) for 10 min at 37°C followed by inactivation with astrocyte media: DMEM (Gibco), 10% fetal bovine serum (Gibco), 1% pen-Strep (Gibco), 1% GlutaMAX (Gibco). The supernatant was quickly removed and replaced with fresh astrocyte media and 200 Kunitz units/mL of DNase. Tissue homogenized using a P1000 pipette, followed by centrifugation at 400 g for 5 min at 4°C. Cells were counted, resuspended in astrocyte media, and plated in T75 flasks (3 pups per flask). 90% of media was changed the day after and subsequently 50% of the media was changed twice a week. After 7-10 days, when cells reached 70% confluency, 3 μM cytosine-arabinoside (AraC; C1768, Sigma-Aldrich) was added to eliminate microglia. To ensure no microglia were left, the flasks were also shaken overnight with 50 mM HEPES added to the media in an orbital shaker incubator (37°C, 100 revolutions per min). The flasks were then rinsed and 0.25% Trypsin-EDTA was added at 37°C for 10 min to lift the astrocytes. After inactivation, the cell suspension was centrifuged at 400 g for 5 min at 4°C, counted and re-plated at a density of 1-2 million cells per well in a 6-well plate. The cells were grown for 1-2 days before further assays. Cells were maintained in an incubator at 37°C and 5% CO_2_.

### Astrocyte conditioned media (ACM) collection and validation

We collected WT and *Mfge8* KO ACM and immunodepleted (ID) MFG-E8 from the WT ACM to obtain three different ACM conditions: WT, MFG-E8 KO ACM and WT MFG-E8 ID and we validated the presence or absence of MFG-E8 in the ACM by ELISA and western-blotting (**Fig S4)**. ACM was collected either from WT or MFGE8 KO astrocyte cultures. The wells were washed with PBS and astrocyte FBS-free media was added overnight. The following day, the ACM was collected for further assays. To immunodeplete MFG-E8 from WT ACM, magnetic dynabeads® M-270 Epoxy (#14301, Thermofisher) were used following the manufacturer’s instructions. MFG-E8 antibody (goat, AF2805, RnD) was coupled overnight to the Dynabeads after which successful coupling was tested by western-blotting with secondary anti-goat HRP of the flow through and input antibody. As a control, an irrelevant antibody was used of the same species and clonality as MFG-E8. Following successful coupling, 1.5 mg of antibody-coated beads were added to ACM and incubated in a rotator for 2 hr. A magnet was used to collect the beads at the tube wall and the supernatant was collected as the immunodepleted ACM. Subsequent ELISA was performed to assess MFG-E8 depletion. The beads were washed with 20 mM HEPES pH 7.5, 150 mM NaCl, 0.1% NP-40 and x1 Halt™ Protease Inhibitor Cocktail (Thermo Scientific, 78429). The immunoprecipitate was eluted in 2% SDS buffer and boiled at 70°C for 10 min. To quantify the levels of MFG-E8 in the ACM, the MFG- E8 quantikine ELISA kit (#MFGE80, RnD systems) was used following the manufacturer’s instructions. ACM from different conditions was assayed in triplicates, along with base astrocyte media as a control. The absorbance of samples was read at 450 nm and wavelength correction was set at 540 nm. For the analysis, the mean absorbance of base astrocyte media and mean absorbance at 0 ng/mL were subtracted. The concentration of MFG-E8 was extrapolated from a standard curve following the manufacturer’s description.

### Synaptosome preparation and conjugation to pHrodo

Synaptosomes were prepared from fresh mouse tissue as previously described using a technique which has been validated by electron microscopy^8,9,105^. 3-5 WT mice aged 2-4 mo were used. The hippocampi and cortices were dissected on ice. Tissue was weighed and homogenized in 5 volumes of sucrose homogenization buffer (5 mM HEPES pH 7.4, 320 mM sucrose,1 mM EDTA, x1 Halt™ Protease Inhibitor Cocktail (Thermo Scientific, 78429) using a Dounce homogenizer with 15-20 strokes. The total homogenate was centrifuged at 3,000 g for 10 min at 4°C. The supernatant was collected and centrifuged again at 14,000 g for 12 min at 4°C. The pellet was carefully resuspended in Krebs-Ringer buffer (KRB: 10 mM HEPES, pH 7.4, 140 mM NaCl, 5 mM KCl, 5 mM glucose, 1 mM EDTA, x1 Halt™ Protease Inhibitor Cocktail (Thermo Scientific, 78429), 45% isotonic percoll solution. The solution was mixed by gently inverting the tube and an interface was slowly created with KRB. After centrifugation at 14,000 g for 2 min at 4°C, the synaptosomal fraction was recovered at the surface of the flotation gradient and carefully re-suspended in KRB to wash. The synaptosomal preparation was centrifuged at again and the pellet was re-suspended in 30-50 uL of KRB. After isolation, a standard BCA protein assay was performed to obtain the amount of protein for subsequent assays.

Synaptosomes were conjugated to low background pH-sensitive dyes, which fluoresce brightly upon acidification (pH 4-6) in late endosomes and lysosomes using an adapted protocol^106^. pHrodo™ Red, succinimidyl ester (P36600) was dissolved as described in the manual. 1 mg of synaptosomes were left at RT on the nutator for 2 hr in sodium bicarbonate 0.1 M in 1 mg/mL pHrodo. After conjugation, synaptosomes were centrifuged at 14,000 g for 1 min, then washed with 1 mL PBS and centrifuged again at 14,000 g for 1 min. Synaptosomes were then resuspended as described by the manufacturer. Next, a standard BCA protein assay on pHrodo conjugated synaptosomes to quantify protein concentrations for subsequent engulfment assays. Synaptosomes were then aliquoted respectively and stored at -80°C.

### *In vitro* microglial synaptosome engulfment assay

The live cell microglial engulfment assay was performed using primary WT or MFG-E8 KO microglia. Microglial media was replaced with either WT, WT MFG-E8 immunodepleted or MFG-E8 KO ACM, 1 hour prior to adding 1 μg of pHrodo-conjugated synaptosomes in the media. Live cell imaging was performed in a cell discoverer 7 (CD7) at 37°C and 5% CO_2_ conditions. Images were captured at 594 nm fluorescence, as well as brightfield (oblique and phase), using x20 magnification (x0.5) at 3-5-min intervals. Cells were imaged for 15 hr, with t=0 being the moment when synaptosomes were added. Analysis was conducted on ImageJ, using the ‘plot z file’ function, following 1 pixel background subtraction pHrodo. Fluorescence intensity at t=0 was subtracted from all intensity values. Results were plotted as pHrodo fluorescence intensity over time and as area under curve (AUC).

### RNA isolation, reverse transcription, and RT-qPCR

For RNA extraction from primary astrocytes or microglia, the cells were lysed and scraped off using TRIzol reagent (15596026, Invitrogen) after which chloroform was added to separate the homogenate layers. RNA was precipitated from the aqueous layer using 2-propanolol and then washed with ethanol. The RNA pellet was re-suspended in nucleus-free water. For RNA extraction from flow cytometry sorted astrocytes or microglia, between 20000 and 30000 cells were collected in RLT lysis buffer with β-mercaptoethanol (1/100). RNA was extracted using the RNAeasy Plus Micro (Quiagen, 74034) protocol following the manufactures description. RNA purity and concentration was assessed by Nanodrop. mRNA was converted to cDNA using the qScript cDNA SuperMix reverse transcription kit as described by the manufacturer (95048, Quantabio). For RT-qPCR, 12ng of cDNA was loaded in triplicates per well in SYBR green PCR master mix (4309155, ThermoFisher) and 200 nM of forward and reverse primers. The reaction was run using a LightCycler 96 Instrument (Roche) with white 96-well plates (04729692001, Roche). Triplicate Ct values were averaged, and data is shown as respective to housekeeping genes (*Actb*, *Gapdh*, *Rpl32*) using the Ct delta method (2^-ΔΔ**Ct**^).

The following primer sequences were purchased from IDT or Merck:

*Actb:* 5’ CATTGCTGACAGGATGCAGAAGG, 3’ TGCTGGAAGGTGGACAGTGAGG

*Gapdh*: 5’ CATCACTGCCACCCAGAAGACTG, 3’ ATGCCAGTGAGCTTCCCGTTCAG

*Rpl32*: 5’ ATCAGGCACCAGTCAGACCGAT, 3’ GTTGCTCCCATAACCGATGTTGG

*Gfap*: 5’ CACCTACAGGAAATTGCTGGAGG, 3’ CCACGATGTTCCTCTTGAGGTG

*Mfge8*: 5 GAAAGCGGTGGAGACAAGGA’, 3’ GCTCAGAACATCCGTGCAAC

*Slc1a2*: 5’ TTCCAAGCCTGGATCACTGCTC, 3’ GGACGAATCTGGTCACACGCTT

*Slc1a3*: 5’ ACCAAAAGCAACGGAGAAGAG, 3’ GGCATTCCGAAACAGGTAACTC

*Aldh1l1*: 5’ CTTCATAGGCGGCGAGTTTGTG, 3’ CGCCTTGTCAACATCACTCACC

*Itgam*: 5’ ATGGACGCTGATGGCAATACC, 3; TCCCCATTCACGTCTCCCA

*Tmem119*: 5’ CCTACTCTGTGTCACTCCCG, 3’ CACGTACTGCCGGAAGAAATC

*Tgfbr1*: 5’ TGCTCCAAACCACAGAGTAGGC, 3’ CCCAGAACACTAAGCCCATTGC

*Cx3cr1*: 5’ GAGTATGACGATTCTGCTGAGG, 3’ CAGACCGAACGTGAAGACGAG

*Mag*: 5’ GGCCGAGGAGCAAGAATGG, 3’ CATGCACTCTGCGATACGCT

*Map2*: 5’ ATGACAGGCAAGTCGGTGAAG, 3’ CATCTCGGCCCTTTGGACTG

*Tubb3*: 5’ TAGACCCCAGCGGCAACTAT, 3’ GTTCCAGGTTCCAAGTCCACC

### Stereotaxic surgeries

Mice were anaesthetized with 4% inhaled Isoflurane (Forane, Abbott Laboratories) and placed in a stereotaxic apparatus (504926, World Precision Instruments Ltd). Anesthesia was maintained at 1.5% in 1.5 L/min oxygen flow. Under aseptic conditions, a midline incision was made to reveal the skull and marcaine (0.025%) was applied locally. Holes were drilled in the skull using a 0.8 mm diameter burr (503599, OmniDrill35 Micro Drill, World Precision Instruments Ltd). Infusion was performed with a Microinjection Syringe Pump (World Precision Instruments Ltd) using a 10 µl Hamilton syringe (NanoFil, World Precision Instruments Ltd) with a fine borosilicate glass capillary (Hamilton). The needle was retracted slowly to avoid backflow 5 min after injection. The incision on the scalp was closed with 6/0 ethicon vicryl suture (W9500T) and Vetbond tissue adhesive (3 M). Subcutaneous carprofen (Carprieve, 5 mg/g body weight) and buprenorphine (Vetergesic, 0.1 mg/g body weight) diluted in 0.9% saline were administered peri-operatively. Animals received carprofen (33.33 mg/mL) in their drinking water for 3 days post-surgery.

**For intracerebroventricular injections of Aβ oligomers**, Aβ 40-S26C synthetic humanized dimers were purchased from Phoenix Pharmaceutical (018-71). 4 uL of either 1 ng/µL Aβ dimers or sterile PBS vehicle were unilaterally injected at a rate 350 nL/min of using the following coordinates: -0.5 mm anterior/posterior, +/-1.0 mm lateral, and -2.3 mm dorsal/ventral from bregma (Paxinos and Franklin’s The Mouse Brain in Stereotaxic Coordinates, Fourth Edition). Mice were harvested either 18 hr or 72 hr post-surgery for microglia-synapse engulfment and synapse loss studies.

**For intrahippocampal injections of tdTomato**, pZac2.1 gfaABC1D-tdTomato was a gift from Baljit Khakh (Addgene viral prep # 44332-AAV5) and was purchased from Addgene at a titer of ≥ 7×10¹² vg/mL. 0.6 uL were unilaterally injected at a rate of 150 nL/min using the following coordinates: -2.0 mm anterior/posterior, -1.2 mm lateral, and -1.65 mm dorsal/ventral from bregma. Mice were harvested 1-month post-surgery for astrocyte morphology studies.

**For intrahippocampal injections of CRISPR-saCas9 constructs**, the following combination of viruses were used: AAV5 *GfaABC_1_D* saCas9-HA alone, AAV5 *GfaABC_1_D* saCas9-HA with AAV5 *U6* control-gRNA *GfaABC_1_D* mCherry or AAV5 *GfaABC_1_D* saCas9-HA with AAV5 *U6 Mfge8*-gRNA *GfaABC_1_D* mCherry. All AAV titers were adjusted to 1.7 x 10^13^ genome copies/mL with sterile saline. A total of 0.6 uL were injected unilaterally at a rate of 150 nL/min using the following coordinates: -2.0 mm anterior/posterior, +/-1.2 mm lateral, and -1.65 mm dorsal/ventral from bregma. hAPP J20 Tg and hAPP NL-F KI mice were harvested at 6 and 8 weeks after respectively for subsequent assays.

### Immunohistochemistry (IHC)

30 μm free-floating tissue sections were washed in PBS followed by pre-treatment in 1% Triton X-100 in PBS for 30 min and rinsed in 0.3% Triton X-100. If a primary mouse antibody was used, M.O.M.® (Mouse on Mouse) Blocking Reagent (Vectorlabs, MKB-2213-1) was added to the pre-treatment. Sections were then blocked in 10-20% serum (donkey or goat), 1% BSA, and 0.3% Triton X-100 in PBS for 2 hr at RT followed by primary antibody incubation overnight at 4°C. Sections were washed in PBS for 10 min, 0.3% Triton X-100 in PBS for 30 min, followed by secondary antibody incubation for 2-3 hr at RT. Sections were then washed in PBS for 10 min, incubated in 1:10000 DAPI in PBS for 10 min and washed in 0.3% Triton for 15 min. 20- month-old mice were treated with an autofluorescence quenching kit (Vector, SP-8400-15), prior to DAPI staining following the manufacturer’s instructions. Finally, sections were mounted onto slides with ProLong Glass mounting medium.

To quantify MFG-E8 protein levels in the hippocampus vs the dorsal striatum, serum and Tx were omitted from the protocol and instead 5% BSA in TBS was used as the buffer base, as Tx washes out lipid-based molecules including MFG-E8^53,100^. To quantify MFG-E8 levels within bulbous vs bushy astrocytes, the standard IHC protocol outlined above was used.

### Immunocytochemistry

Cells were fixed for 10 min at RT in 4% PFA, 4% sucrose in PBS, pH 7.4 followed by treatment with 0.25% Triton X-100 in PBS for 3 min at RT. Blocking of cells was performed for 1 hr at RT in 3% serum, 1% BSA. Cells were then incubated in primary antibodies overnight at 4 °C. Next, the cells were washed in PBS and incubated with secondary antibodies for 1 hr at RT. After a final 10 min wash with PBS, cells were incubated in 1:10,000 DAPI in PBS for 10 min.

### *In situ* hybridization (RNAScope)

RNAScope was performed following the manufacturer’s instructions using the RNAscope Fluorescent Multiplex Assay (ACDBio, 320293). The following probes were purchased from the manufacture: *Slc1a2*-C1 (441341), *Cx3cr1*-C1 (314221), *Mfge8*-C2 (408771), *Aldh1l1*-C3 (405891), *Itgav*-C3 (513901), *Itgb3*-C1 (481451), *Itgb5*-C2 (404311). 12 μm thick sections were mounted on Superfrost Plus GOLD Slides (Thermo Fisher Scientific, K5800AMNZ72). The slides were incubated in H_2_O_2_ for 4 min at RT and washed in RNAse-free water. Slides were then placed in boiling target antigen retrieval for 4 min, dehydrated in 100% ethanol for 5 min and treated with Protease Plus for 15 min at RT before probe hybridization for 2 hr at 40°C. For IHC staining post-RNAScope, the slides were blocked in 2% serum 0.01% Triton X- 100 for 30 min and then incubated with the primary antibodies (1/100) overnight at 4°C. The following day, the slides were washed and incubated with the secondary antibodies for 2 hr at RT. The blocking buffer was used to dilute the antibodies. Slides were then incubated with DAPI and mounted onto slides with ProLong Glass mounting medium.

### *In vivo* microglial engulfment imaging and analysis

10-15 μm z-stack images were acquired at an interval of 0.3 μm on a x63 magnification Zeiss LSM 800 confocal microscope. Four to six regions of interest in the hippocampal CA1 stratum radiatum per hemisphere were acquired to include 6-9 cells per mouse or experimental replicate. Imaris 3D surface rendering was used to create a surface of either the P2Y12 or GFAP/tdTomato channel. Within that surface, either CD68 or Lamp1 was masked, and a further surface was then created using a volume filter of > 0.01 μm^3^. Finally, Homer1 was masked within the lysosomal surface and a further surface was created using a volume filter of > 0.001 μm^3^. The engulfment index is quantified using the following formula: (volume of engulfed material in lysosome /cell surface volume) x100.

### Super-resolution Airyscan imaging and analysis

Images were acquired on a Zeiss LSM 880 microscope with Airyscan detector using a 63x, 1.4NA oil immersion Plan-Apochromat objective. A zoom factor of 1.8x and frame size of 2048x2048 was used for all images, resulting in an XY pixel size of 37 nm. Z-step size was 144 nm, with 8 steps per Z-stack, resulting in a stack thickness of 1.15 µm. Three regions of interest were acquired in the hippocampal CA1 stratum radiatum per hemisphere for each experimental replicate. Super-resolution synapse images were processed in Zen Black using 3D Airyscan Processing at strength 6.0. Synapse quantification was performed using Imaris software. The spots detection function was used to generate spots, with Region Growing, Shortest Distance Calculation and Background Subtraction enabled. Spots size was selected to maximize the detection of immunoreactive puncta (XY diameter 0.15 µm, Z diameter 0.45 µm). Spots were classified using the ‘Intensity Center’ filter. Local contrast was used to define the spot growth boundary, using a value that appropriately reflected the immunoreactive signal of the source channel. A volume filter was then applied to remove spots smaller than 0.005 μm^3^. Spots were colocalized using a MATLAB script (Colocalize Spots XTension) at a 0.25 μm distance.

### Post-mortem human tissue immunofluorescence

Post-mortem human formalin-fixed paraffin-embedded tissue was obtained from the Newcastle Brain Tissue Resource. All tissue samples were donated with full and informed consent and were obtained under the appropriate ethical and material transfer approval. The cohort included pathologically diagnosed AD cases and neurologically normal controls. The level of AD pathology in all cases was assessed using current diagnostic consensus criteria. Accompanying clinical and demographic data of all cases used in this study can be found in **Table 1**.

**Table 1.** Human post-mortem tissue demographics.

| Case | Disease category | Age | Sex | Braak LB Stage | Braak Tangle Stage | PM Delay (h) | Thal | TREM2 mutation | Apoe polymorphism |
| --- | --- | --- | --- | --- | --- | --- | --- | --- | --- |
| 1 | AD | 81 | Male | Amygdala-only | 6 | 41 | 5 | None | 3/4 |
| 2 | AD | 87 | Male | 0 | 6 | 21 | 5 | None | 3/4 |
| 3 | AD | 86 | Male | 0 | 6 | 9 | 5 | None | 3/4 |
| 4 | AD | 85 | Female | 0 | 6 | 21 | 5 | None |  |
| 5 | AD | 84 | Male | 0 | 6 | 24 | 5 | None |  |
| 6 | Control | 88 | Male | 0 | 2 | 28 | 1 | None | 2/3 |
| 7 | Control | 94 | Male | 0 | 3 | 25 | 1 | None | 3/4 |
| 8 | Control | 91 | Male | 0 | 3 | 24 | 0 | None | 3/4 |
| 9 | Control | 88 | Female | 0 | 2 | 22 | 2 | None |  |
| 10 | Control | 87 | Male | 0 | 2 | 72 | 3 | None |  |

8 µm tissue sections were mounted on adhesive slides. The sections were incubated in a 60°C oven overnight before staining after which they were deparaffinized in xylene and rehydrated in decreasing grades of alcohol. For heat-induced antigen retrieval, the slides were boiled in 0.1 M citrate buffer (pH 6.0) in a pressure cooker for 10-15 min. The sections were blocked in 10% donkey serum, 1% BSA, and 0.1% Triton X-100 in TBS for 30 min at RT and then incubated with the primary antibodies overnight. The sections were washed in 0.1% Triton X- 100 TBS and incubated with the secondary antibodies for 1 hr at RT. The sections were then washed in 0.1% Triton X-100 TBS, treated with an autofluorescence quenching kit (Vector, SP-8400-15) following the manufacturer’s instructions, incubated in 1:10000 DAPI in PBS for 10 min, washed again and mounted onto slides with ProLong Glass mounting medium. For MFG-E8 IHCs, the antigen retrieval step was replaced with 1 μg/mL proteinase K (Promega, PRMC5005) digestion in buffer (6.1g Trizma Base/ Tris, 0.11 g CaCl_2_ in 1 L H_2_O, pH 7.6) at 37°C for 30 min followed by boiling in a pressure cooker for 10 min in 10mM EDTA (pH 6.0) and the primary antibodies were left for 48 hr at RT.

### *Mfge8* sgRNA design and *in vitro* screening

The sgRNA sequences were designed using the CHOPCHOP website. sgRNA sequences targeting exons 1 or 2 were preferred to avoid truncated protein production. Sequences were selected depending on predicted efficiency (> 0.6), location (exons 1 and 2), GC content (between 40-70%) and low predicted mismatch. DNA templates containing the sgRNA nucleotide sequences were designed for transcription. The PAM sequence was removed from the sgRNA sequence, and the following sequence was added at the 5’ end for recognition of the template by T7 RNA polymerase during transcription:

TTCTAATACGACTCACTATA.

The following sequence was added at the 3’ end of the sgRNA nucleotide sequence to facilitate the recognition of the sgRNA by the saCas9:

GTTTTAGTACTCTGGAAACAGAATCTACTAAAACAAGGCAAAATGCCGTGTTTATCTCGTCAACTTGTTGGCGAGATTT.

A complementary oligonucleotide sequence of the final product was also designed to create a double stranded DNA template via annealing. The DNA template was annealed by a PCR reaction with the forward and complementary oligonucleotides at 10 μM, with Q5® High-Fidelity DNA Polymerase (#M0491, NEB) according to the manufacturer’s protocol. The double stranded DNA template was used for *in vitro* transcription using the HiScribe™ T7 Quick High Yield RNA Synthesis Kit (#E2050, NEB), as described by the manufacturer. For transcription, 1 μg of DNA template was loaded in the reaction. DNase treatment was performed to remove the DNA template according to the manufacturer. RNA was subsequently purified using the Monarch RNA Cleanup Kit (50 µg) (#T2040, NEB) according to the manufacturer’s instructions. Quantification and purity assessment of RNA was performed using a Nanodrop. Amplification of the appropriate genomic DNA (gDNA) regions using the previously designed primers, was conducted using 20 ng of mouse gDNA (#G3091, Promega) and the Q5® High-Fidelity DNA Polymerase. Finally, an *in vitro* digestion was performed to assess the cleavage efficiency of the sgRNA molecules. A PCR reaction was performed using 100 nM of the purified RNA product, 200 ng of the amplified gDNA region product and 1 μM EnGen® Sau Cas9 (#M0654, NEB), according to the manufacturer, and running the reaction at 37°C for 1 hr, followed by 5 min at 75°C. The final products were visualized via gel electrophoresis using 2-3% Agarose (Invitrogen) gels. Agarose gels were prepared by boiling 2-3% agarose in TBE and adding Gel Red (41003, Biotium) at 1:10,000 concentration. Gel loading dye (#B7025, NEB) was added at 1:6 concentration to each sample and the Quick-Load® Purple 100 bp DNA Ladder (#N0551, NEB) was used as a size indicator. Gels ran at 100V in 1x TBE running buffer for 1-1.5 hr and were visualized in the Amersham Imager 680 (Bioke) system. Three successful sgRNA sequences were selected, cloned into plasmids, and packaged into AAV5 viral constructs commercially to yield the following construct: AAV5 (*U6 Mfge8*gRNA x3) *GfaABC_1_D* mCherry **(Fig S9)**.

### Cell isolation for flow cytometry and sort

Fresh cortex and hippocampi were processed using the adult brain dissociation kit (Miltenyi Biotec, 130-107-677). After enzymatic digestion and homogenization, the cell suspension was filtered through a 70 μM cell strainer before mixing with debris removal solution. Cells were centrifuged at 300 g for 10 min, washed with ice-cold FACS buffer (PBS, 2% FBS, 0.78 mM EDTA) and incubated for 30 min at 4 °C with FACS buffer containing Fc block (BD Biosciences) and primary antibody mix. Experiments were performed on a BD Aria III (BD Biosciences). Dead cells were excluded using DAPI stains. Microglia population was gated based on CD45^+^CD11b^+^CX3CR1^+^CD206^-^ **(Fig S5)**. Astrocytes were gating based on CD45^-^ ACSA2^+^ **(Fig S5)**. Flow cytometry data were analyzed using FACSDiva software 4.0 and FlowJo 10 software (Treestar).

### Western blotting

A BCA protein assay was used to determine the amount of protein in fractions from the synaptosome preparation. 40 μg of protein were loaded on 4-12% Bis-Tris gels (Thermofisher Scientific) and run with appropriate sample buffer and running buffer. Gels were transferred to nitrocellulose membranes using an iBlot 2 Dry Blotting System (Thermofisher Scientific) as described by the manufacturer. Blots were blocked in casein PBS (1:1, Biorad) for 30 min at RT on shaker and then probed overnight at 4°C on shaker with primary antibodies (1:1,000) in casein PBS-T (0.01%). Blots were washed then probed with secondary HRP-antibodies in casein PBS-T (0.01%) for 1 hr at RT on shaker, washed, treated with Immobilon Forte western HRP substrate (Millipore, WBLUF0500) and then visualized on an Amersham Imager 680 (Bioke) system. Blots were stripped with x1 ReBlot Plus Strong Antibody Stripping Solution (Millipore, 2504), re-blocked and re-probed as outlined above.

## Antibodies

For mouse immunostaining, the following primary antibodies were used: rabbit anti-mouse IBA1 (Wako Chemicals, 019-19741; 1/500), rat anti-mouse CD68 (Bio-Rad, MCA1957; Clone FA-11, 1/200), rabbit anti-mouse P2Y12 (Anaspec, AS-55043A; 1/500), chicken anti-mouse Homer1 (Synaptic System, 160 006; 1/200), mouse anti-mouse 6E10 (Biolegend, 803001; Clone 6E10, 1/200), rabbit anti-mouse Bassoon (Synaptic System, 141 003; 1/200), mouse anti-mouse Synaptophysin (Abcam, ab8049, 1/200), goat anti-mouse MFG-E8 (RnD, AF2805, 1/200), rabbit anti-mouse GFAP (Merck, AB5804, 1/1000), chicken anti-mouse GFAP (Abcam, ab4674, 1/1000), guinea pig anti-RFP (Synaptic Systems, 390 004, 1/500), mouse anti-HA (Biolegend, 901514, 1/200), rabbit anti-mouse p62/SQSTM1 (Abcam, ab91526, 1/2000), rat-anti-mouse LAMP1 (Abcam, Ab25245, 1/200), guinea pig anti-rat S100β (Synaptic Systems, 287-004, 1/200), rabbit anti-mouse S100β (Abcam, ab41548, 1/200), rabbit anti-rat EAAT1 (Abcam, Ab416, 1/200), rabbit anti-mouse NeuN (Abcam, ab104225, 1/1000), rabbit anti-mouse Ezrin (Cell Signalling, 3145S, 1/200).

For human immunostaining, the following primary antibodies were used: rabbit anti-human SQSTM1/p62 (Abcam, ab91526, 1/200), goat anti-human S100β (R&D Systems, AF1820, 1/200), mouse anti-human ALDH1L1 (Merck, MABN495, 1/200), mouse anti-human MFG-E8 (NovusBio, NBP2-44412-0, 1/100), mouse anti-human MFG-E8 (OriGene, BM607S, 1/100), mouse anti-human MFG-E8 (Abcam, AB17787, 1/100), mouse anti-human MFG-E8 (RnD Systems, MAB27671, 1/100), rabbit anti-human GFAP (Abcam, AB68428,1/200).

Secondary antibodies used were a combination of Alexa Fluor 488, 594 and 647 (1/500, Jackson ImmunoResearch and Thermo Fisher Scientific) from either donkey or goat where appropriate.

For western blotting, the following antibodies were used: rabbit anti-mouse GAPDH (Abcam, ab181602, 1/20,000), goat anti-rabbit HRP (Abcam, ab205718, 1/10,000), hamster anti-mouse MFG-E8 (Origene, AM26517AF-N, 1/1000) and rabbit anti Armenian hamster (Abcam, Ab5745, 1/1000).

For flow cytometry, the following antibodies were used: BUV395 CD45 (BD Biosciences, 564279; 1/400), Pe-Cy7 CD11b (BD Biosciences, 552850; 1/400), BV421 CX3CR1 (Biolegend, 149023; 1/400), PE ACSA-2 Antibody, anti-mouse (Miltenyi Biotec, 130-123-284)

### Image acquisition

Most images were acquired either on a Zeiss LSM800 confocal microscope (×20 objective 0.8-NA, ×40 objective 1.3NA oil and 63×0.8-NA oil) or a Leica Stellaris 8 STED confocal microscope (×20 objective 0.8-NA, x40 objective 1.3NA oil and 100×0.8-NA oil). Some images were also acquired using an Axio Scan Z1 slide scanner (x40 objective) and a Cell discoverer 7. The same settings including laser intensity and pixel size were kept constant for all sections in the same immunostaining. Typically, a frame size of 1024 x 1024 pixels was used for most experiments. The appropriate step size and stack size was determined based on the experiment: Low magnification with very large field of view: 1-2.5 µm step size and 30-60 µm stack size (e.g., supplementary NeuN staining for CRISPR studies of both hippocampi and cortices), medium magnification with large/medium field of view: 0.3-0.5 µm step size and 30- 50 µm stack size (e.g., whole hippocampal hemisphere images to count number of bulbous astrocytes), high magnification with medium/small field of view: 0.2-0.3 µm step size and 10- 30 µm stack size (most data acquired in the manuscript) or 8-12 µm stack size (for RNAScope and human immunostaining studies). Acquired data was analyzed either on ImageJ Fiji or Imaris software where appropriate. Representative images are obtained either from ImageJ Fiji or Imaris and are adjusted for brightness, contrast, and background where appropriate.

### Image J Fiji analysis

The software ImageJ Fiji was used to perform analysis of 2D data shown in µm^2^. Background subtraction, despeckle, thresholding, and masking were used where appropriate. Thresholding analysis was performed on a 2D maximum intensity projection of the Z-stack. Data was analyzed either for the whole imaged ROI or further ROIs were selected using the ROI manager e.g., for astrocyte area single astrocytes were chosen from a larger ROI. The channels were split, and an appropriate threshold was applied using dark background. To identify regions of colocalization between two or more channels e.g., mCherry^+^ GFAP^+^ cells, the image calculator function AND was used. The analyze particles function was used for quantification, and if necessary, an appropriate a size exclusion criteria was applied e.g., for DAPI nuclei > 5 µm or cells > 10 µm. This resulted in a total count, total area, average size and % of area for the particles in each channel. For the Iba1 Sholl analysis, two concentric circles were drawn on ImageJ using the at 40 µm or 80 µm away from the center of the astrocyte and the number of Iba1 cells was manually quantified. The number of bulbous astrocytes in a region of interest was manually quantified either through the stack or as a 2D maximum Z-projection, depending on the size of the Z-stack, using the ROI function as an identifier. Representative images have been adjusted for brightness and contrast.

### Imaris analysis

The software Imaris was used to perform analysis of 3D data shown in µm^3^. Data was analyzed using the surface and spots functions. To identify and or quantify the number of cells/specific cell type, a 3D rendered surface was made for DAPI and the staining of a cell surface marker e.g., GFAP/NeuN/S100β or a cell specific gene e.g., *Cx3cr1*/*Slc1a2* was masked ‘within’ the DAPI surface, and a further surface was created of the masked channel. The ‘quality’ filter was used to adjust the area and volume of surfaces according to fluorescence signal. The function ‘separate touching objects’ was enabled, and the count of different cell types was automatically modelled by Imaris. To quantify the number of mRNA puncta, the staining of a gene of interest e.g., *Mfge8* was masked ‘within’ the cell identifying surface and the puncta number was counted using the spots function. The diameter and number of spots were adjusted depending on the fluorescence signal using the ‘quality’ filter. To quantify the volume of MFG-E8 within bulbous vs bushy astrocyte processes, a surface was made of the S100β immunostaining and the MFG-E8 immunostaining was masked within. A surface was made for the internalized MFG-E8, and the volume was calculated per 1500 μm^3^. To quantify the volume of Ezrin within bulbous vs bushy astrocyte processes, a surface was made of the S100β immunostaining, and the Ezrin immunostaining was 3D rendered using the spots function. The surface-spot colocalization function was used to analyze the Ezrin spots at a 0 μm distance from the S100β surface. Finally, the total volume of the Ezrin spots on the S100β surface was divided by the S100β cell volume. To quantify the distance between 6E10-immunoreactive plaques p62-accumulated bulbous astrocytes, a surface was made for both, and the shortest distance calculation was used. The data acquired from the surface and spots functions resulted in a total count and total volume as shown in the figures. Representative images have been adjusted for brightness and contrast.

### Single cell RNA sequencing data presentation

Re-presentation of Linnarsson lab single cell RNA sequencing database^54^ **(Fig S6E)** was conducted using R (version 4.2.0).

### Statistics

All statistical analyses were performed in Prism (GraphPad Software, Version 10.3.0, July 26, 2024) as appropriate. Statistical methods were based on similar previously published experiments. Experimenters were blinded to the genotype and treatment of the mice as well as the demographics of the human post-mortem tissue. Outliers were identified and removed from the dataset using GraphPad Prism (ROUT, Q=1%). For datasets where multiple ROIs were sampled per mouse, outlier analysis was done per ROI before calculating the mouse average. Normal distribution and equality of variance of the residuals were then tested per mouse average using the Shapiro-Wilk normality test and the F-test, Spearman’s test or Bartlett’s test, using significance level α= 0.05. To stabilize variances on heteroscedastic residuals, we performed Y=log(Y) transformations before fitting the two-way ANOVA model (stated in the figure legends where applicable). All statistical tests were performed per mouse average or culture average to avoid pseudoreplication. Statistical significance was set at α = 0.05. To compare two groups, either a two-tailed unpaired student’s t-test, two-tailed unpaired student’s t-test with a Welch’s correction, two-tailed paired student’s t-test, two-tailed Mann-Whitney tests or Wilcoxon matched pairs signed rank test was used depending on nature of the data. To compare three groups, either a one-way ANOVA with Bonferroni’s multiple comparisons post-hoc test, Brown-Forsythe and Welch ANOVA with Dunnett’s T3 multiple comparisons test or Kruskal Wallis with Dunn’s multiple comparisons test was used. To compare more than three groups, a two-way ANOVA with Bonferroni’s multiple comparison post hoc, multiple Mann-Whitney tests followed by the Bonferroni-Dunn correction, Wilcoxon matched-pairs signed rank test followed by Bonferroni-Dunn correction or multiple unpaired or paired t-tests with Bonferroni-Dunn correction. All statistical tests and data representation are clearly stated in the figure legends. Where applicable interaction term is provided as (Interaction: F (DFn, DFd), p-value). All data shown as mean ± SEM. p-values shown ns P>0.05; *P<0.05; **P<0.01; ***P<0.001; ****P<0.0001.

